# Coexistence of piRNA and KZFP defense systems: Evolutionary dynamics of layered defense against transposable elements

**DOI:** 10.1101/2025.11.27.690821

**Authors:** Yusuke Nabeka, Hideki Innan

**Author notes:** Corresponding author: SOKENDAI, Research Center for Integrative Evolutionary Science, The Graduate University for Advanced Studies, Hayama, Kanagawa 240-0193, Japan. (Y.N.); (H.I.).

## Abstract

Transposable elements (TEs) pose a persistent threat to genome stability, and host organisms have consequently evolved sophisticated defense mechanisms to restrain them. In animals, the most prominent systems include PIWI-interacting RNAs (piRNAs) and Krüppel-associated box zinc-finger proteins (KZFPs). Because both systems recognize TEs in a sequence-specific manner and induce epigenetic silencing, they appear functionally redundant at first glance. However, KZFPs are a relatively recent innovation that emerged and diversified in genomes where the piRNA pathway was already established. This raises an important question: under what conditions can a second, seemingly redundant defense system invade and persist? To address this, we constructed a mathematical model integrating the evolutionary dynamics of TEs, piRNAs, and KZFPs. Our approach focuses on a key mechanistic asymmetry between the two defense systems: whereas piRNA-mediated suppression is dependent on TE activity, KZFPs provide constitutive suppression that does not rely on ongoing TE activity. We show that these distinct modes of action generate interactions that extend beyond simple redundancy or additivity. We derive analytical conditions under which KZFPs can invade a pre-existing TE-piRNA equilibrium and characterize the evolutionary logic that enables stable coexistence of these multilayered defense strategies. Together, our results provide a theoretical framework for understanding how complex, layered genome defense systems can evolve and persist.

## Introduction

Transposable elements (TEs) are mobile genetic elements capable of moving and replicating within host genomes, posing a significant threat to genomic stability (Kazazian 2004). Uncontrolled proliferation of TEs can lead to gene disruption and chromosomal rearrangements, which, in extreme cases, may result in the extinction of entire host populations. Consequently, hosts have evolved multiple defense mechanisms to suppress TE activity.

Among these suppression mechanisms, epigenetic regulation that silences TE transcription is a prominent strategy. Two well-characterized examples are the PIWI-interacting RNA (piRNA) pathway and Krüppel-associated box (KRAB) zinc-finger proteins (KZFPs) (Lawlor and Ellison 2023). piRNAs are small RNAs present in most eukaryotes and are derived from specific genomic loci known as piRNA clusters (Ozata *et al*. 2019). When a TE inserts into a piRNA cluster, the host generates piRNAs complementary to the TE sequence. These piRNAs bind to PIWI clade proteins, targeting TEs through sequence complementarity. The piRNA/PIWI complex recruits histone methyltransferases (HMTs) and DNA methyltransferases (DNMTs), which catalyze histone methylation (H3K9me3) and DNA methylation. These epigenetic modifications induce heterochromatin formation around the TE insertion site, effectively suppressing TE transcription (Brennecke *et al*. 2007; Halic and Moazed 2009; Cosby *et al*. 2019; Iwasaki *et al*. 2025).

In contrast, KZFPs represent a more recent evolutionary innovation, found in coelacanths, lungfish, and tetrapods, originating approximately 420 million years ago (Huntley *et al*. 2006; Tadepally *et al*. 2008; Emerson and Thomas 2009; Iwasaki *et al*. 2025). KZFPs consist of a DNA-binding zinc-finger array and a KRAB domain. The zinc fingers recognize specific DNA triplets, enabling sequence-specific binding to TEs, while the KRAB domain recruits TRIM28 (also known as KAP1), DNMTs, and other chromatin modifiers to establish repressive heterochromatin around the targeted TE locus, robustly suppressing its transcription (Emerson and Thomas 2009; Levin and Moran 2011; Ecco *et al*. 2017; Yang *et al*. 2017; Imbeault *et al*. 2017; Cosby *et al*. 2019; Almeida *et al*. 2022; Rosspopoff and Trono 2023; de Tribolet-Hardy *et al*. 2023; Iwasaki *et al*. 2025).

While both piRNAs and KZFPs suppress TE activity through sequence-specific recognition and epigenetic modifications, it remains unclear why KZFPs evolved in the presence of the already effective piRNA pathway. One might hypothesize that KZFPs operate more efficiently than the piRNA pathway; however, this does not fully explain their coexistence, because less effective defense systems are generally expected to be lost over evolutionary time. Conversely, if their suppressive effects were merely additive, one might expect the hosts to accumulate as many defense mechanisms as possible. The reality—that these two systems coexist, while new, similar systems have not arisen indefinitely—presents an intriguing question. Here, we focus specifically on their roles in TE suppression in order to isolate the evolutionary logic of how apparently redundant TE-defense systems can emerge and coexist, while we acknowledge that defense systems are sometimes strongly coopted for essential roles outside TE suppression, such as developmental processes, and their loss could be lethal (Mani *et al*. 2014; Hanin *et al*. 2026).

A clue to resolving the question of why the apparently redundant defense systems coexist may lie in the different mechanisms upon which their functions depend. The piRNA system requires TE transposition, as piRNAs are generated from TE insertions within piRNA clusters (Brennecke *et al*. 2007; Kofler 2019; Tomar *et al*. 2023; Srivastav *et al*. 2024). Although it relies on stochastic insertion events, it substantially decreases the transposition rate once established. In contrast, KZFPs can evolve recognition specificity over longer timescales by accumulating mutations in their DNA-binding zinc-finger arrays (Nardelli *et al*. 1992; Looman *et al*. 2002; Ecco *et al*. 2017; Najafabadi *et al*. 2017; Zhang *et al*. 2022; Wells *et al*. 2023). Once evolved, their persistence is determined by a balance between their own suppression efficacy and cost, independent of the TE activity level. In other words, a newly emerged KZFP is less specific at first, and its silencing effect may be modest until the binding array is refined. It is these asymmetries that may give rise to interactions between the two systems that go beyond simple exclusion or additivity.

We thus hypothesize that these mechanistic differences in TE dependence and suppressive specificity are the key to the coexistence of the two TE suppression mechanisms: the two distinct TE suppression systems are neither mutually exclusive nor function in a simple additive manner, but rather interact with each other. To test this hypothesis, we construct a mathematical model that describes the dynamical interactions among TEs, piRNAs, and KZFPs, reflecting their different modes of TE suppression. Through the analysis of this model, we elucidate the evolutionary logic of multilayered genome defense strategies by clarifying (1) the conditions under which KZFPs can invade and become established in an existing TE-piRNA equilibrium, (2) the conditions under which both systems can stably coexist, and (3) the consequences of their coexistence for the host’s TE copy number and fitness.

Previous theoretical work has examined the population dynamics and regulation of TE copy number, including the effects of selection, self-regulation of transposition, and host modifiers (Charlesworth and Charlesworth 1983; Charlesworth and Langley 1986). Recent studies have also highlighted evolutionary interactions between TEs and piRNAs (Kofler 2019; Luo *et al*. 2020; Tomar *et al*. 2023). However, these studies do not explain why an additional KZFP system should evolve alongside the piRNA system. This gap motivates this study.

## Model

Our objective in this study is to explain why seemingly redundant TE suppression mechanisms are maintained within genomes and to determine the evolutionary conditions for their emergence and coexistence. To this end, we have constructed a mathematical model that integrates the evolutionary dynamics of TE copy number, piRNA-producing alleles, and KZFP alleles.

Our model is based heavily on the “trap model” of piRNA-mediated suppression described by Tomar *et al*. (2023), which assumes that a piRNA-producing allele completely suppresses transposition. We extend their work by explicitly incorporating the evolutionary dynamics of KZFP-mediated TE suppression. Tomar et al. (2023) considered three versions of the trap model: (i) a neutral model, in which neither TEs nor piRNA-producing alleles affect host fitness; (ii) a deleterious TE–neutral cluster model, in which TE insertions are deleterious but piRNA-producing alleles are neutral; and (iii) a deleterious TE–deleterious cluster model, in which both TE insertions and piRNA-producing alleles carry fitness costs. Of these three cases, we focus exclusively on model (iii), because only this case is relevant to the long-term invasion and maintenance of KZFP-mediated suppression. In model (i), KZFP is not favored because TEs impose no fitness cost and there is no selective advantage to suppressing them. In model (ii), active TEs are eventually eliminated by piRNA-mediated suppression, thereby eliminating the selective pressure required to maintain KZFP-mediated suppression.

We consider a deterministic model of an infinite diploid population where genetic drift can be ignored. We also assume that the population is always in Hardy-Weinberg equilibrium. The genome of each individual contains an infinite number of potential TE insertion sites, together with a single piRNA cluster locus and a single KZFP locus. For simplicity, we consider a single TE family whose sequence does not mutate and we assume free recombination so that all TE insertion sites, the piRNA-cluster locus, and the KZFP locus are well mixed in the population. We assume the piRNA locus has two allelic states: a piRNA-producing allele that fully suppresses TEs, and a null allele with no suppressive function. Similarly, the KZFP locus has two allelic states: a KZFP suppressor allele with partial suppression efficacy ranging from 0 to 1, and a null allele with no suppressive function. Under these assumptions, we construct the system of three coupled ordinary differential equations, describing (i) the change in mean TE copy number, (ii) the frequency of a piRNA-producing allele and (iii) the frequency of a KZFP suppressor allele. The detailed derivations of (i), (ii) and (iii) are given below.

### Change in TE Copy Number

First, we derive an equation for the dynamics of the mean TE copy number, which is governed by the balance between gain via transposition and loss due to selection. We define *n* as the population mean number of TE copies located outside of the piRNA clusters. Since new transpositions arise from existing TE copies, the expected increase in mean TE copy number per generation is proportional to the current mean TE copy number *n*, scaled by the basal transposition rate per copy *u*, giving *un*. This transposition is suppressed by both piRNA and KZFP systems and we assume their effects are multiplicative. The fraction of individuals lacking piRNA-mediated suppression (i.e., homozygous for non-piRNA alleles) is (1− *p*)^2^, assuming Hardy-Weinberg equilibrium and dominance of the piRNA-producing allele. The average suppression effect of KZFPs is 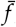, so the fraction of events escaping KZFP suppression is 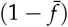. The derivations of *p* and 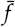 are detailed in the following sections. Then, the effective transposition rate per copy is 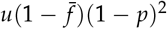 and the corresponding total increase in TE copy number per generation is 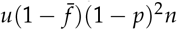. Since *n* counts TE copies located outside the piRNA clusters, this increase should strictly be multiplied by (1 −*π*), where *π* denotes the piRNA cluster fraction, defined as the fraction of the genome occupied by piRNA clusters. Here, we approximate 1− *π* ≈1 because *π* is very small and its direct contribution to the total TE copy number is negligible; for example, piRNA clusters occupy only approximately 0.1–5% of typical metazoan genomes (Rosenkranz *et al*. 2022).

For the decrease of *n* via selection, we assume that each TE copy reduces the host fitness by a selection coefficient of *s*. We assume that the fitness of an individual carrying *n*_*i*_ TE copies is given by 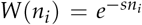 and the corresponding Malthusian fitness is 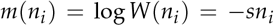 . Treating the individual TE copy number (*n*_*i*_) as a quantitative trait, the change in the population mean TE copy number *n* due to selection in continuous time can be approximated by 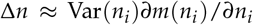 . Assuming random mating with free recombination, the TE copy number is approximately Poisson-distributed, so Var(*n*_*i*_) *n* (Charlesworth and Charlesworth 1983). Because *∂m*(*n*_*i*_)/*∂n*_*i*_ = −*s*, the change in *n* due to selection is approximated as −*sn*, which is a standard approximation in quantitative genetic models of the dynamics of TE copy number. It should be noted that Charlesworth and Charlesworth (1983) pointed out that such a linear relationship between Malthusian fitness and the TE copy number is insufficient to maintain a stable equilibrium of TE copy number in the absence of self-regulation of TE activity. In contrast, in the recent theoretical work on the piRNA trap model, Tomar *et al*. (2023) demonstrated that the linear model is indeed sufficient for stability, because the piRNA system itself provides the necessary regulatory feedback. We therefore adopt this linear approximation, which allows the derivation of explicit theoretical predictions. Combining the terms derived above gives the full expression for the change in TE copy number:

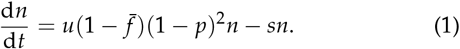

### Change in piRNA Allele Frequency

We next consider the dynamics of the piRNA allele frequency *p*, which is determined by direct generation via TE insertions and selection. We assume the piRNA cluster locus has two allelic states: a non-functional, null allele (frequency 1 − *p*) and a functional, piRNA-producing allele (frequency *p*). The change in *p* is driven by three factors: direct generation through TE insertions into the piRNA cluster, the benefit arising from the reduction in new deleterious TE insertions, and the cost of the cluster insertion. Therefore, we assume that 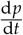 involves a direct gain term, a benefit term, and a cost term.

The direct gain term is derived from the rate at which non-functional, null alleles are converted into functional, piRNA-producing ones. This process requires two conditions: a TE transposes, and it lands within the piRNA cluster. As described above, the effective transposition rate per copy is 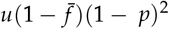. A fraction *π* of these transposition events will occur within the piRNA cluster, potentially creating a new functional, piRNA-producing allele. Therefore, the rate of these insertion events is 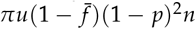. To translate this rate of allele creation into a rate of frequency change in a diploid population, we must correct for the total number of alleles at this locus in the population (two per individual). This introduces a scaling factor of 1/2. Thus, the rate of increase in the frequency *p*, or the direct gain term, is given by

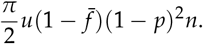

For the benefit and cost terms, we use standard weak selection approximations for allele frequency change in continuous time (Crow and Kimura 1970, p. 192). The benefit term represents the selective advantage of piRNA-producing alleles, which arises from their ability to reduce the copy number of deleterious TEs outside the piRNA clusters. In a genotype lacking piRNA-mediated suppression, the expected number of new TE insertions per generation is 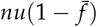. Since each TE copy carries a fitness cost *s*, the selective advantage of carrying a piRNA-producing allele, *B*_*π*_, is given by

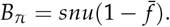

Because piRNA-mediated suppression is assumed to be dominant, this advantage is expressed in both heterozygotes and homozygotes carrying the piRNA-producing allele. Under weak selection for a dominant advantageous allele, the contribution of this selective advantage to the change in *p* is given by

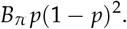

The cost term follows from selection against an additive deleterious allele:

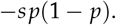

Here, we assume that the piRNA-producing allele itself contains one deleterious TE insertion within the piRNA cluster and that its effect is additive. Thus, the allele incurs a fitness cost equivalent to carrying one TE copy with selection coefficient *s*. This cost is accounted for separately from the main TE copy number described in Equation (1), because *n* denotes TE copies outside the piRNA cluster.

Combining the direct gain term, the benefit term and the cost term, the dynamics of *p* are given by

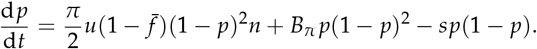

Substituting 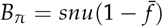, we obtain

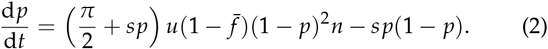

### Change in KZFP Allele Frequency

We derive the dynamics of the KZFP suppressor allele frequency by focusing on its net selective advantage. KZFPs are proteins that exert their suppressive effect by binding to TE sequences via a DNA recognition motif. Unlike the piRNA system, this mechanism does not have a direct gain process where a TE is required to be inserted into a specific locus to trigger a response. Instead, we assume that the suppression efficacy is determined by the sequence similarity between the KZFP’s recognition motif and the target TE copy. For simplicity in this study, we do not explicitly model the recognition sequence itself; instead, we treat the suppression efficacy as a trait value. We focus here on a minimal two-allele model for analytical tractability and clarity, while a more general formulation allowing an infinite number of KZFP alleles with varying suppression efficacies is given in Appendix A. Specifically, we consider a functional suppressor allele with frequency *x* and suppression efficacy *f* (0 ≤ *f* ≤ 1), and a non-functional null allele with frequency 1 − *x* and no suppressive function ( *f* = 0). Assuming that the suppressor allele is dominant over the null allele with respect to its suppressive function, the average suppression efficacy across the population under Hardy-Weinberg equilibrium is:

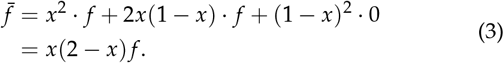

We focus on a short evolutionary timescale after the invasion of the KZFP suppressor allele, during which mutation between the suppressor allele and the null allele can be ignored. The change in the frequency of the suppressor allele over time is therefore determined by natural selection:

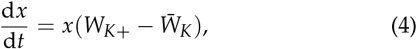

where *W*_*K*+_ is the average fitness of individuals carrying at least one suppressor allele, and 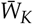 is the population mean fitness. The two fitness terms will be derived below.

We define the net selective advantage of the suppressor allele as *σ*, which is given by the benefit of TE suppression minus the maintenance cost. The benefit arises from the reduction in the number of new deleterious TE insertions. This can be quantified by comparing the rate of transposition experienced by individuals with and without the suppressor allele. For individuals without the suppressor allele, the rate of new TE insertions is *nu*(1 −*p*)^2^. For individuals carrying at least one suppressor allele, the transposition rate is reduced by a fraction *f*, resulting in a rate of *nu*(1− *f*)(1 −*p*)^2^. The difference between these two rates represents the number of TE insertions prevented per generation by carrying the suppressor allele, which is *nu f* (1− *p*)^2^. Since each TE copy carries a fitness cost of *s*, the total benefit conferred by the suppressor allele is this reduction in TE copy numbers multiplied by *s*. We also assume a constant maintenance cost of *c* for carrying the suppressor allele, which is independent of its suppression efficacy (the null allele is assumed to have no cost). Therefore, the net selective advantage of the suppressor allele, *σ*, is given by

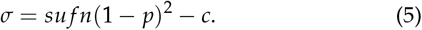

Under the weak selection approximation for a dominant allele with the net selective advantage *σ*, the population mean fitness is 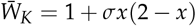 and the average fitness of individuals carrying at least one suppressor allele is *W*_*K*+_ = 1 + *σ*. Substituting 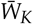 and *W*_*K*+_ into Equation (4), we obtain

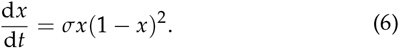

The three differential equations (1), (2) and (6) provide the entire theoretical basis used in the following analyses.

## Results

In this article, we analyze the long-term outcomes after a second defense mechanism, the KZFP system, is introduced into a stable TE-piRNA equilibrium, which we refer to as the “pre-invasion” state. Our analysis proceeds as follows: First, we derive the analytical conditions that determine whether a rare KZFP allele can invade in this pre-invasion state. We then characterize the two possible long-term outcomes of a successful invasion: either the KZFP allele spreads to fixation in the population (see below for definition) or it is maintained at a polymorphic equilibrium.

### Conditions for KZFP Invasion

This work considers the invasion of a KZFP allele into a stable system composed only of TEs and piRNAs. To do so, we need to first investigate the conditions for the TE-piRNA preinvasion equilibrium before the KZFP invades. The TE-piRNA pre-invasion equilibrium can be explored by setting the KZFP frequency to zero (*x* = 0) to reduce the three-variable model to a two-variable dynamical system with TE copy number (*n*) and piRNA allele frequency (*p*):

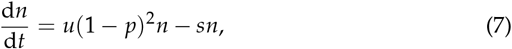

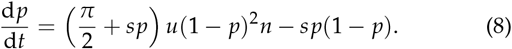

A nontrivial equilibrium (*n*^∗^, *p*^∗^) of this system is given analytically by solving 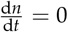 and 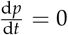:

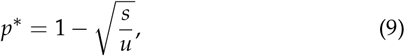

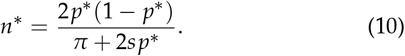

This equilibrium requires *s* < *u* for both TEs and piRNAs to persist. At this equilibrium, *p*^∗^ is determined solely by the ratio *s*/*u*, whereas *n*^∗^ additionally depends on *π*. This equilibrium is locally stable, as shown in Appendix B.

Given this equilibrium, we are interested in whether the KZFP system can invade into the population. In practice, in our deterministic treatment, we investigate the net selective advantage of the KZFP suppressor allele when it is very rare (i.e, *x* ≈ 0). From Equation (6), 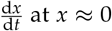is approximately given by

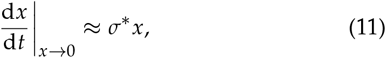

where the initial net selective advantage, *σ*^∗^, is

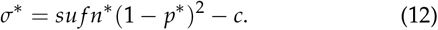

For a KZFP allele to successfully invade, the initial net selective advantage *σ*^∗^ must be positive, meaning the benefit from TE suppression must exceed its maintenance cost. By substituting the pre-invasion equilibrium values of *n*^∗^ and *p*^∗^, given by Equations (9) and (10), into Equation (12), we have the condition where the invasion of KZFP is possible:

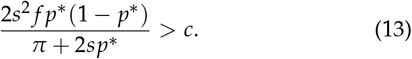

Figure 1 illustrates this condition in the ( *f, c*) parameter plane for various values of *s, u*, and *π*. The colored curves represent the boundaries separating regions where KZFP invasion is possible (below the curves, Equation (13)) from those where it is not (above the curves). We refer to these boundaries as the invasion thresholds. As shown in Figure 1A, increasing *s* shifts the invasion threshold upward, unless *s* is unrealistically large. This means that a large *s* allows KZFPs to invade even when their maintenance cost *c* is high. This is because a TE with a large *s* imposes a strong deleterious effect on the host fitness so that suppressing TEs is beneficial, thereby making such KZFPs likely to invade.

**Figure 1.**
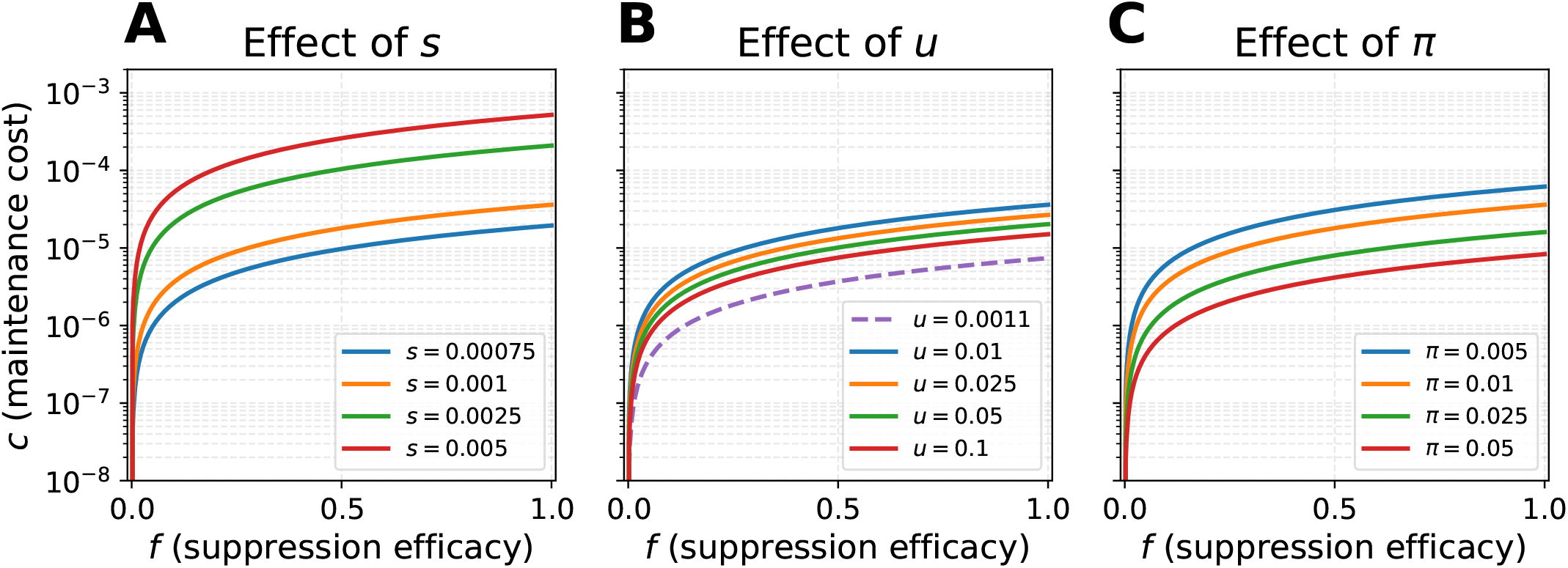
Effect of the parameters on the KZFP invasion threshold. Each curve represents the analytical invasion threshold derived from Equation (13), plotted in the suppression efficacy ( *f*) versus maintenance cost (*c*) parameter space. KZFP invasion is possible in the region below each respective curve. (A) Effect of varying the selection coefficient, *s*. (B) Effect of varying the basal transposition rate, *u*. (C) Effect of varying the piRNA cluster fraction, *π*. Unless otherwise specified in the panel legend, the fixed parameters are *s* = 0.001, *u* = 0.01, and *π* = 0.01.

The effects of the other two parameters, *u* and *π*, are also investigated in Figure 1B and C. Figure 1B shows that a large *u* value allows only KZFP alleles with low cost to invade, except when *u* is very small. This can be explained as follows: when transposition is highly active (i.e., *u* is large), the piRNA pathway is already strongly engaged in TE suppression, resulting in a high *p*^∗^ and low *n*^∗^ (Equations (9) and (10)). In this state, an additional benefit of KZFP invasion is limited. As a result, KZFP invasion is only possible when the cost *c* is sufficiently low. An exception is when *u* is very small, as indicated by the purple dashed line in Figure 1B. In such a case, transposition activity is very low and TEs are already rare, making the benefit of KZFP is small. As a result, only KZFP alleles with very low cost can invade. Likewise, Figure 1C shows that large *π* values also restrict the invasion to KZFPs with low cost. This is because a large piRNA cluster enhances the efficiency of the piRNA pathway, leading to a reduction in *n*^∗^ (Equation (10)), which diminishes the benefit of KZFP.

### Characterization of Possible Long-Term Outcomes after KZFP Invasion

To understand possible long-term outcomes, we consider the equilibrium after successful invasion. This equilibrium is referred to as the “post-invasion equilibrium”. Let (*n*_eq_, *p*_eq_, *x*_eq_) denote the equilibrium values of (*n, p, x*) at the post-invasion equilibrium. These values were obtained by numerically integrating Equations (1),(2), and (6) until the system converged to equilibrium. Figure 2 shows the post-invasion equilibrium values of (*n, p, x*) across the ( *f, c*) parameter space for four different values of the selection coefficient *s* (*s* = 5 × 10^−3^, 2.5 ×10^−3^, 1× 10^−3^, and 7.5 × 10^−4^), with *u* = 0.01 and *π* = 0.01 fixed. The red curves correspond to the invasion thresholds derived in the previous section. The post-invasion equilibria are locally stable, as shown in Appendix B.

**Figure 2.**
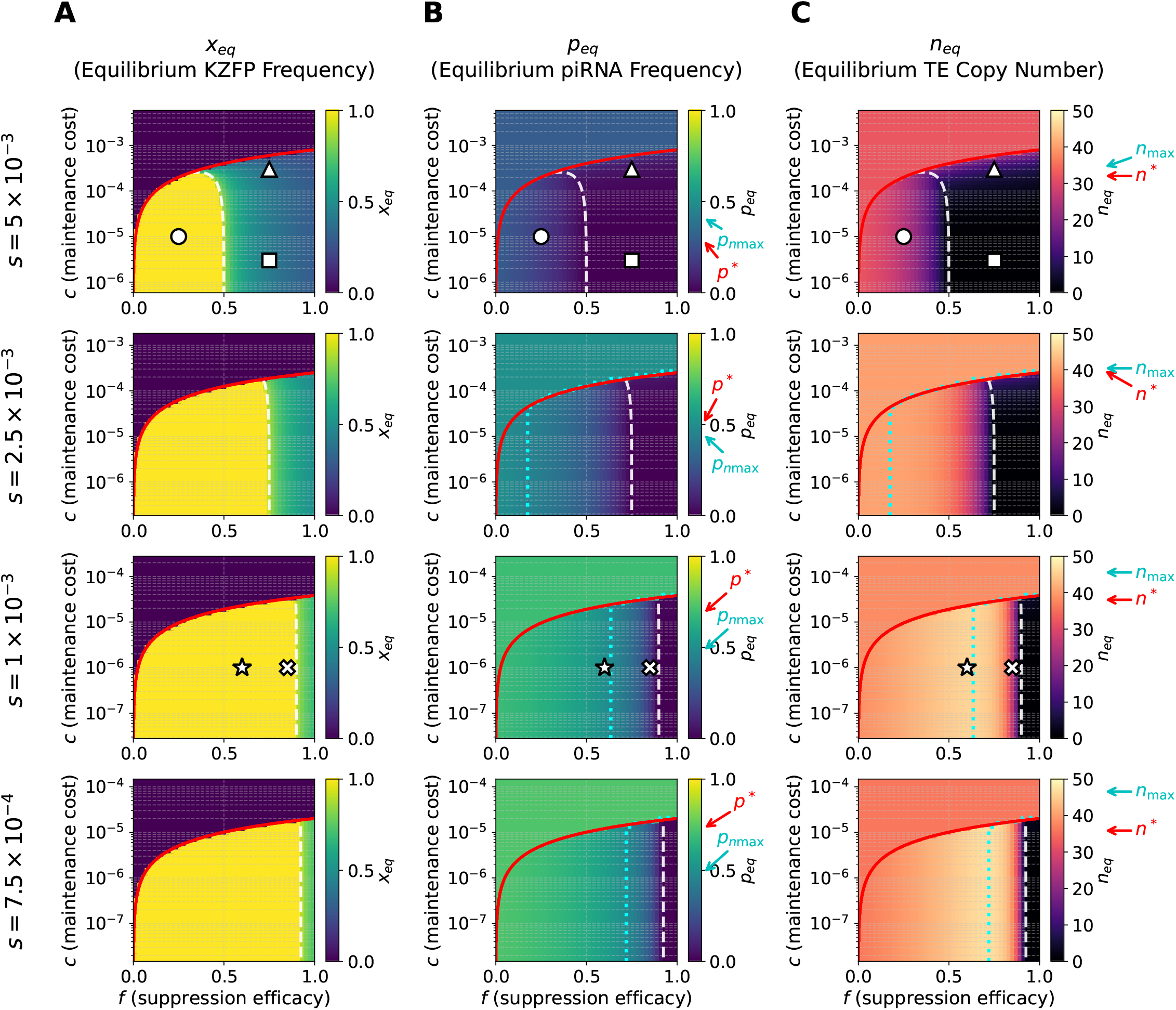
Post-invasion equilibrium values in the ( *f, c*) parameter space. (A) Equilibrium frequency of the KZFP allele (*x*_eq_), (B) equilibrium frequency of the piRNA-producing allele (*p*_eq_), and (C) equilibrium TE copy number (*n*_eq_) after KZFP invasion plotted in the parameter space of suppression efficacy of KZFP ( *f*) and maintenance cost of KZFP (*c*). Four different values of the selection coefficient *s* are used, while *u* = 0.01 and *π* = 0.01 are fixed. The post-invasion equilibrium values are numerically computed by using the solve_ivp ODE solver in the scipy package (Virtanen *et al*. 2020). The color bars are to present the value of focal quantities in the post-invasion equilibrium. The red curve shows the invasion threshold from Equation (13), below which KZFP invasion is possible. The white dashed curve shows the upper boundary of the KZFP-fixation zone from Equation (22). The cyan dotted line in the lower two panels of (B) and (C) marks ( *f, c*) at which the piRNA frequency (*p*_*n*max_) maximizes the equilibrium TE copy number, *n* = *n*_max_ (see Equation (27)). The cyan arrows on the color bars indicate *p*_*n*max_ and *n*_max_, together with the pre-invasion equilibria *p*^∗^ and *n*^∗^ with red arrows. Symbols mark the parameter sets whose dynamics are shown in Figure 5: circle (Figure 5A; *s* = 5 × 10^−3^, *f* = 0.25, *c* = 1 × 10^−5^); cross (Figure 5B; *s* = 1 × 10^−3^, *f* = 0.85, *c* = 1 × 10^−6^); star (Figure 5C; *s* = 1 × 10^−3^, *f* = 0.6, *c* = 1 × 10^−6^); triangle (Figure 5D; *s* = 5 × 10^−3^, *f* = 0.75, *c* = 3 × 10^−4^); square (Figure 5E; *s* = 5 × 10^−3^, *f* = 0.75, *c* = 3 × 10^−6^).

Let us focus on *x*_eq_, the frequency of the KZFP allele at the post-invasion equilibrium, which is visualized in Figure 2A. It is clearly shown that *x*_eq_ is almost 1 over a broad region of the parameter space, shown in yellow. We define this parameter region as the “KZFP-fixation zone”. It should be noted that, in our deterministic model, the KZFP allele cannot mathematically reach fixation in the population; however, we consider it effectively fixed when there is a consistent selective pressure driving its frequency toward 1 across all frequencies.

Figure 2A also shows that *x*_eq_ has an intermediate value in the region in green. We define this region as the “KZFP-polymorphic zone”. To explore the boundary between these two zones, we again focus on *σ*, the net selective advantage of the KZFP suppressor allele, which should be given by a function of *x*. As defined, within the KZFP-fixation zone, the net selective advantage should be positive for any value of *x* between 0 and 1, so that the KZFP suppressor allele increases to 1. In contrast, this should not hold in the KZFP-polymorphic zone, resulting in an equilibrium KZFP frequency *x*_eq_ less than 1. A detailed derivation of this boundary is as follows.

To obtain the net selective advantage of KZFP as a function of *x*, we use a quasi-equilibrium approximation in which it is assumed that the variables *n* and *p* immediately reach their quasi-equilibrium value, denoted by (*n*_qs_, *p*_qs_). This approximation is justified by a clear time-scale separation between the fast (*n, p*) dynamics and the slow *x* dynamics (see Appendix C). This reduces the 3D system to an effective 1D dynamics in *x*. Under this assumption, 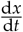 is expressed as

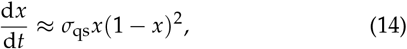

where *σ*_qs_, the net selective advantage of KZFP at the quasi-equilibrium, is given by Equation (5):

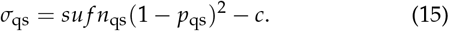

The quasi-equilibrium (*n*_qs_, *p*_qs_) is given by solving 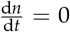 and 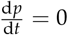 from Equations (1) and (2) with 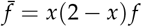:

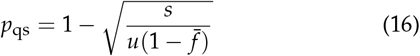

and

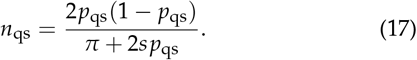

These equations show that the piRNA frequency and TE copy number at the post-invasion equilibrium in the KZFP-fixation zone are independent of the KZFP cost, *c*.

We then obtain the explicit form of the net selective advantage of KZFP, *σ*_qs_, as a function of *p*_qs_, by substituting (*n*_qs_, *p*_qs_) into Equation (15),

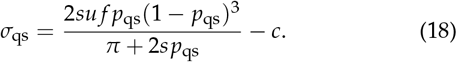

Note that *p*_qs_ is a monotonically decreasing function of *x* in the interval [0, 1]. As *x* varies over 0 ≤ *x* ≤ 1, *p*_qs_ ranges over *p*_*x*1_ ≤ *p*_qs_ ≤ *p*_*x*0_, where

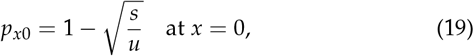

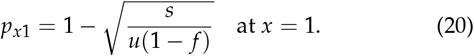

Therefore, we should evaluate *σ*_qs_ only for *p*_qs_ ∈ [*p*_*x*1_, *p*_*x*0_].

Equation (18) is useful for visualizing how the fate of KZFP (fixed or polymorphic) is determined. Figure 3 plots *σ*_qs_ with *p*_qs_ for two representative values of *f* . *σ*_qs_ is a function of *p*_qs_ with *σ*_qs_(*p*_qs_ = 0) = *σ*_qs_(*p*_qs_ = 1) = −*c*. We can only focus on *σ*_qs_ within its biologically feasible range, [*p*_lower_, *p*_upper_], shown by the orange shaded region in Figure 3. Because *p*_qs_ is a frequency, it must satisfy 0 ≤ *p*_qs_ ≤ 1. In addition, because *p*_qs_ is induced by *x* ∈ [0, 1], it must also lie in the interval [*p*_*x*1_, *p*_*x*0_]. Therefore, the biologically feasible range is the intersection of these two intervals, namely

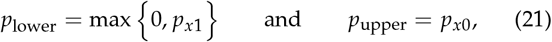

**Figure 3.**
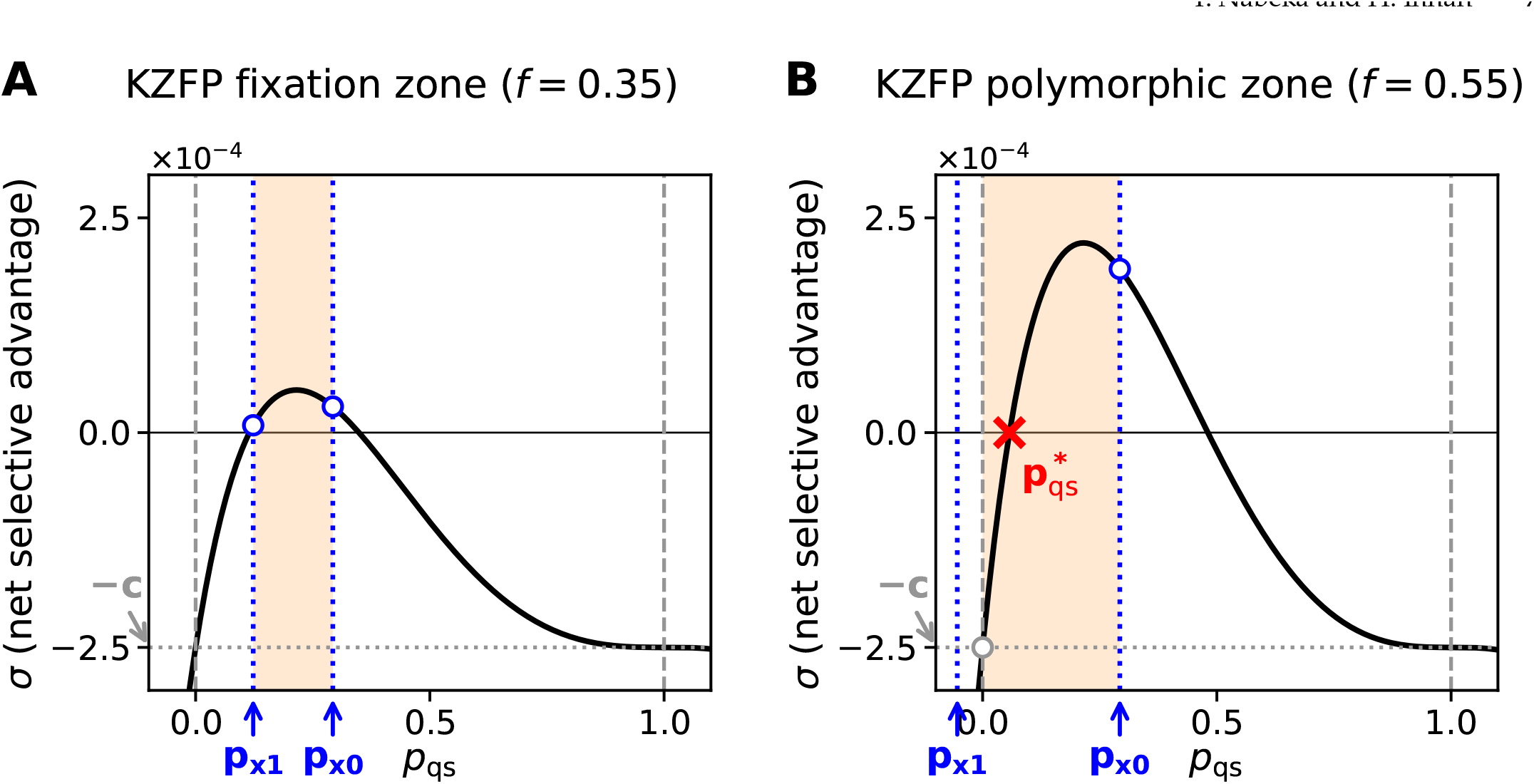
Net selective advantage of the KZFP allele frequency, *σ*_qs_, as a function of the quasi-equilibrium piRNA frequency *p*_qs_ for (A) *f* = 0.35, representing the KZFP-fixation zone, and (B) *f* = 0.55, representing the KZFP-polymorphic zone. The parameters are fixed at *s* = 5 × 10^−3^, *u* = 0.01, *π* = 0.01, and *c* = 2.5 × 10^−4^. The black curve shows *σ*_qs_ as a function of *p*_qs_, given by Equation (18). The gray vertical dashed lines at *p*_qs_ = 0 and *p*_qs_ = 1 delimit the biologically relevant range for 0 ≤ *p*_qs_ ≤ 1 and *n*_qs_ ≥ 0. The blue dotted lines indicate the feasible range of *p*_qs_ corresponding to 0 ≤ *x*≤ 1, from *p*_*x*1_ to *p*_*x*0_. The orange shaded region indicates the feasible range of *p*_qs_, which is the overlap of these constraints. The red cross in (B) shows the point where *σ*_qs_ = 0 at 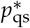 within the feasible range. In (A), *σ*_qs_ is positive throughout the feasible range, so the KZFP allele is driven toward fixation. In (B), *σ*_qs_ changes sign within the feasible range, indicating that the system converges to a stable polymorphic equilibrium with 0 < *x*_eq_ < 1.

where *p*_*x*0_ < 1 under the condition in which pre-invasion equilibrium exists, *s* < *u*. As *p*_qs_ increases from *p*_lower_, *σ*_qs_ increases to a single peak and then declines. If *σ*_qs_ crosses zero inside the feasible range of *p*_qs_, we denote the value of *p*_qs_ at this intersection by 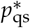 (i.e., 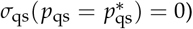.

Figure 3 shows two plots, representing typical cases in the KZFP-fixation and -polymorphic zones. Figure 3A shows a plot for *f* = 0.35, in which *σ*_qs_ is positive in the entire feasible range of *p*_qs_, indicating KZFP can essentially fix. By contrast, Figure 3B illustrates a case in the KZFP-polymorphic zone ( *f* = 0.55). In this plot, *σ*_qs_ is 0 at 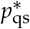, which is inside the feasible range of *p*_qs_. It is indicated that the net selective advantage, *σ*_qs_, is positive for 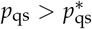, whereas negative for 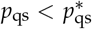. Given this behavior of the net selective advantage, the KZFP frequency reaches a stable equilibrium with an intermediate value, 0 < *x*_eq_ < 1. In other words, the KZFP suppressor allele is polymorphic.

The boundary between the KZFP-fixation and KZFP-polymorphic zones is obtained when *σ*_qs_ is zero in the feasible range, i.e., when 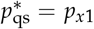. The border is obtained by solving

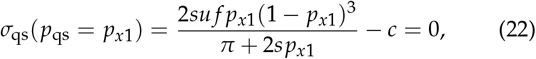

which is shown by the white dashed curve in Figure 2.

### KZFP-fixation zone

We explore the behavior of the frequency of piRNA and TE copy number in the KZFP-fixation zone. Figure 2B shows the equilibrium frequency of piRNA, *p*_eq_, computed by Equation (2). In the KZFP-fixation zone, *p*_eq_ is always lower than *p*^∗^ (from bright to dark colors in Figure 2B). This is intuitively easy to understand. At equilibrium, 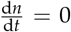 and 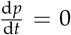 hold simultaneously. From 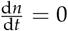 we obtain

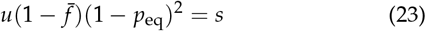

because 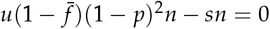, and *n*_eq_ > 0 allows cancellation of *n*, where 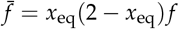 is the average suppression efficacy of KZFP at its equilibrium from Equation (3). In the KZFP-fixation zone, as *x*_eq_ = 1 and 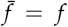 hold, by solving Equation (23) for *p*_eq_, we obtain

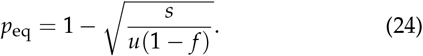

Thus, for fixed *u* and *s*, KZFP with any *f* (> 0) necessarily lowers *p*_eq_ relative to *p*^∗^. It is indicated that the two suppression systems cooperate in the fixation zone, with KZFP almost fixed while the frequency of piRNA decreased.

Figure 2C illustrates the equilibrium TE copy number, *n*_eq_, computed from Equation (1). In the KZFP-fixation zone, *n*_eq_ is typically smaller than its pre-invasion level *n*^∗^, when selection is strong (Figure 2C, the upper panel). However, this does not necessarily hold when selection is weak as shown in the lower three panels in Figure 2C with moderate KZFP suppression efficacy, *f*, where *n*_eq_ exceeds *n*^∗^, which may appear counter-intuitive. The condition for *n*_eq_ > *n*^∗^ could be obtained by expressing *n*_eq_ as a function of *p*_eq_. At equilibrium, 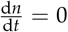 and 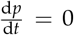 hold simultaneously, therefore we have 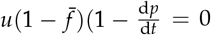. Substituting this into 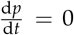, we then obtain the explicit relationship between *n*_eq_ and *p*_eq_ at equilibrium:

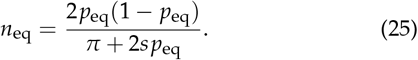

Equation (25) provides a convenient tool for assessing under what condition *n*_eq_ can exceed the pre-invasion level, *n*^∗^. Figure 4 plots the relationship between *n*_eq_ and *p*_eq_ given by Equation (25) for strong and weak selection cases (*s* = 0.005 for the strong case; Figure 4A, *s* = 0.001 for the weak case; Figure 4B). In both cases, *n*_eq_ is unimodal with a single peak at *p*_*n*max_, indicated by the star. Here, we denote the value of *p*_eq_ that maximizes the TE copy number by *p*_*n*max_, where *n*_max_ is the value of *n*_eq_ at the peak (we will derive *p*_*n*max_ and *n*_max_ later). *p*^∗∗^ is the value that produces *n*_eq_ = *n*^∗^ in the left side of the star (orange circle). It is important to note that the feasible range of *n*_eq_ is determined by the feasible range of *p*_eq_ in one-to-one correspondence, and that, within the fixation zone, *p*_eq_ is given by a monotonically decreasing function of *f* (see Equation (24)). This relationship is visualized by showing the corresponding *f* value by colored points along the curve of Equation (25), from *f* = 0 by cyan circle to the border value of *f* by red circle (calculated by solving Equation (22)). In the strong-selection case in Figure 4A, *p*^∗^ < *p*_*n*max_ holds with any *f* value, so that *n*_eq_ is maximized at *f* = 0 and never exceeds *n*^∗^. By contrast, in the weak case in Figure 4B, as *p*^∗^ exceeds *p*_*n*max_, the feasible range of *p*_eq_ contains the star, at which *n*_eq_ has its maximum *n*_max_. This means that, if *f* is low ( *f* <∼ 0.8 so that *p*_eq_ > *p*^∗∗^), *n*_eq_ exceeds *n*^∗^. If *f* is sufficiently high (i.e., *f* > ∼0.8), as well as the strong selection case, *n*_eq_ is smaller than *n*^∗^. Thus, our analytical result indicates *n*_eq_ can exceed *n*^∗^ under certain conditions, although it may be somehow counter-intuitive.

**Figure 4.**
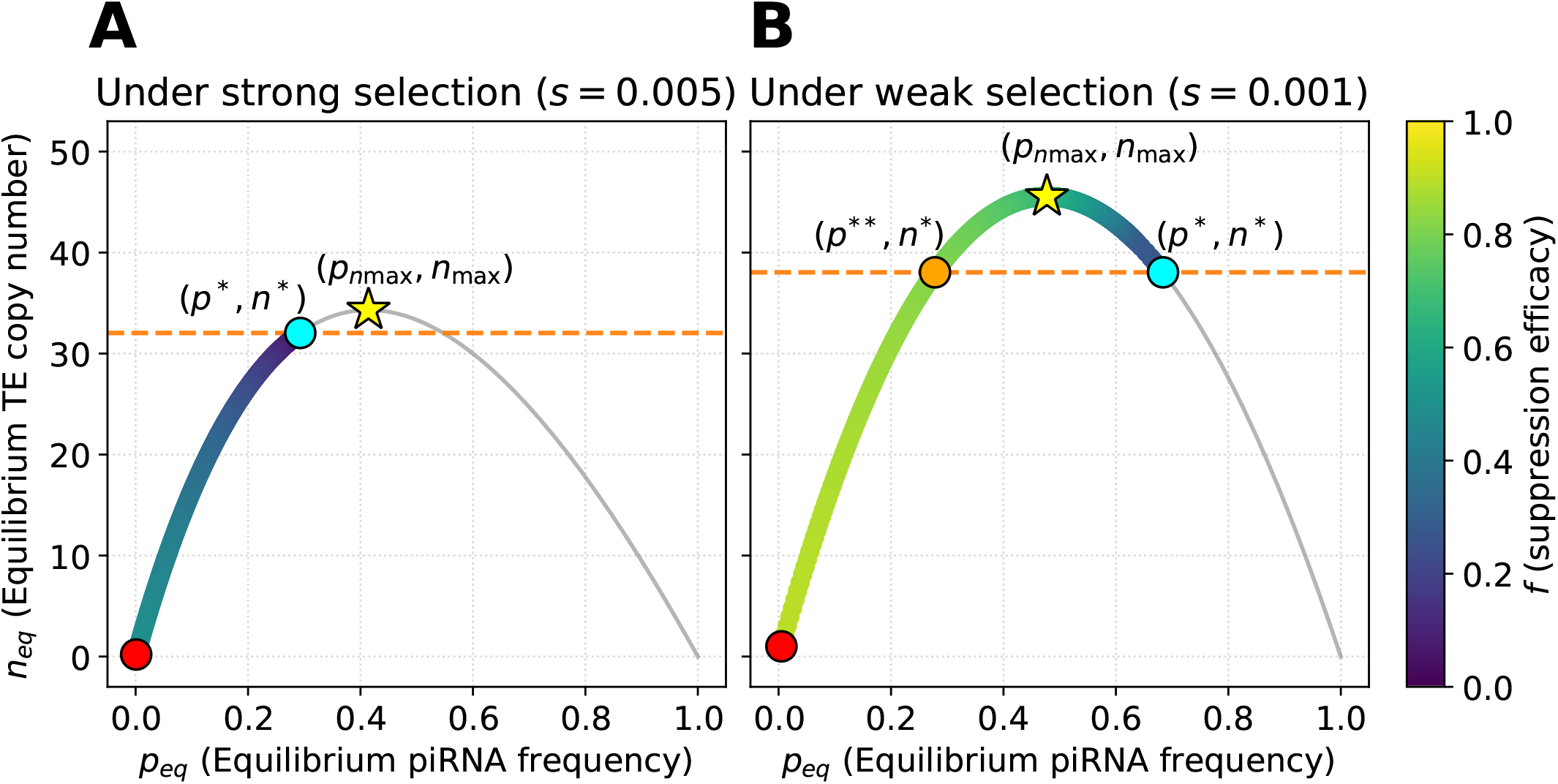
Post-invasion equilibrium relationship between TE copy number and piRNA frequency (*n*_eq_, *p*_eq_) as a function of the KZFP suppression efficacy *f* . Fixed parameters: *u* = 0.01, *π* = 0.01, and *c* = 10^−5^. (A) Strong selection (*s* = 0.005). The pre-invasion value of *p* (*p*^∗^) lies to the left of the peak (*p*^∗^ < *pn*_max_), so all post-invasion equilibria have *n*_eq_ at or below the pre-invasion level *n*^∗^. (B) Weak selection (*s* = 0.001). The pre-invasion value of *p* (*p*^∗^) lies to the right of the peak (*p*^∗^ > *pn*_max_). In the range *p*^∗∗^ < *p*_eq_ < *p*^∗^, *n*_eq_ exceeds the pre-invasion level *n*^∗^. The gray curve shows the analytical equilibrium mapping given by Equation (25). The color bar represents the value of *f* . Cyan circle: pre-invasion equilibrium (*n*^∗^, *p*^∗^) at *f* = 0. Red circle: post-invasion equilibrium at full suppression efficacy ( *f* = 1). Star: local maximum *n*_max_ attained at *pn*_max_ . Orange horizontal line: pre-invasion TE copy number *n*^∗^. Orange circle: point (*n*^∗^, *p*^∗∗^) where *n*_eq_ = *n*^∗^ at *f* ≈ 0.8.

The condition under which *n*_eq_ can exceed *n*^∗^ (i.e., *p*^∗^ > *pn*_max_) can be expressed as a function of *s, u* and *π* as follows. We first derive the piRNA frequency that maximizes TE copy number, *p*_*n*max_. Differentiation of Equation (25) with respect to *p*_eq_ gives

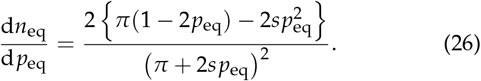

Then, the biologically relevant root of this, denoted by *p*_*n*max_, is obtained exactly as

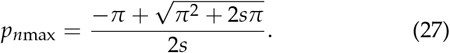

Substituting *p*_*n*max_ into Equation (25), we have a local maximum, denoted by *n*_max_:

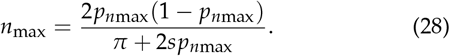

For *n*_eq_ to be greater than *n*^∗^, *p*^∗^ > *p*_*n*max_ has to hold:

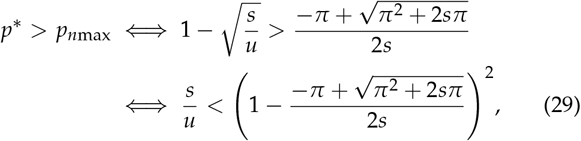

where *p*^∗^ is from Equation (9). For *s*/*π* « 1, this condition can be approximated as

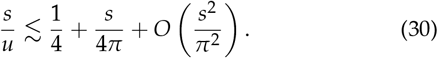

Thus, the ratio *s*/*u*, together with the terms involving *s*/*π*, determines whether the equilibrium TE copy number increases or decreases after KZFP invasion.

In addition to the equilibrium values of these three quantities, *x, p* and *n*, we explore how they change over time after the invasion of KZFP. Figure 5 shows the results for five parameter sets at the locations of the open circle, triangle, square, star, and cross in Figure 2. Here we still focus on the KZFP-fixation zone, that is, the open circle, star, and cross, in Figures 5A–C. When *p*^∗^ < *p*_*n*max_ (i.e., strong selection), Figure 5A shows that *x* rises to fixation while *p* declines moderately (e.g. from ∼0.3 to ∼0.2), consistent with Equation (24). *n* also decreases monotonically. When *p*^∗^ > *p*_*n*max_ (weak selection), as shown in Figures 5B and C, *x* and *p* follow similar dynamics to those in Figure 5A, except that the behavior of *n* is complicated. In Figure 5B with a large *f, n* initially increases above *n*^∗^ (from ∼ 38 to ∼ 45) and then declines to ∼ 29, whereas with intermediate *f* (Figure 5C), *n* increases from ∼ 38 to ∼ 45 and remains elevated at equilibrium.

**Figure 5.**
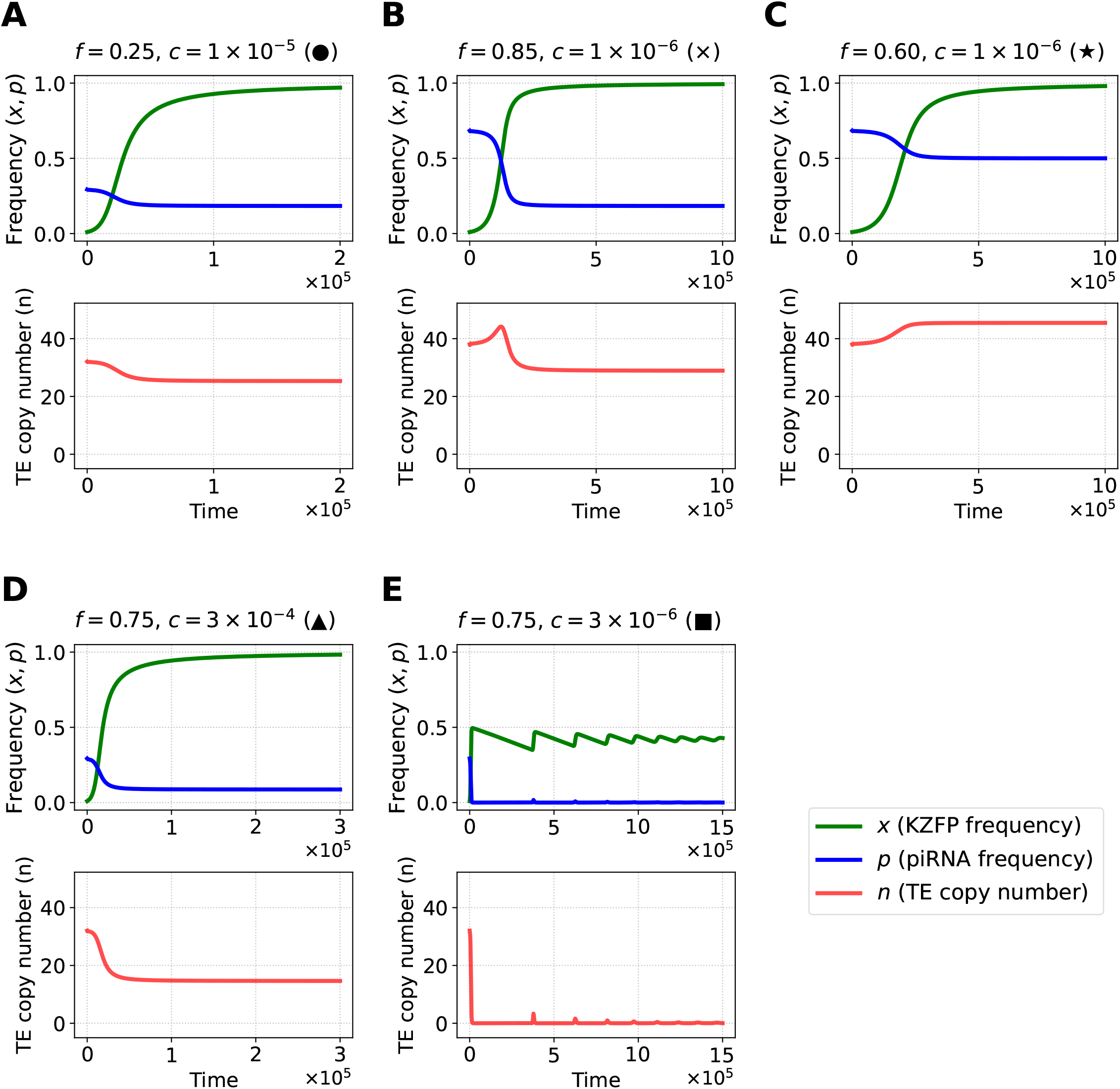
Dynamics of KZFP frequency (*x*), piRNA frequency (*p*), and TE copy number (*n*) for five representative parameter sets sampled from Figure 2. In each column, the upper panel shows the time courses of *x* (green) and *p* (blue), and the lower panel shows the time course of *n* (red). All trajectories start from the pre-invasion equilibrium (*n*^∗^, *p*^∗^), assuming an initial KZFP frequency of *x* = 0.01. (A) KZFP-fixation zone with strong selection (*s* = 5 × 10^−3^, *f* = 0.25, *c* = 1 × 10^−5^; circle in Figure 2). (B) KZFP-fixation zone with weak selection and high suppression efficacy (*s* = 1 × 10^−3^, *f* = 0.85, *c* = 1 × 10^−6^; cross in Figure 2). (C) KZFP-fixation zone with weak selection and moderate suppression efficacy (*s* = 1 × 10^−3^, *f* = 0.6, *c* = 1 × 10^−6^; star in Figure 2). (D) KZFP-polymorphic zone with high maintenance cost (*s* = 5 × 10^−3^, *f* = 0.75, *c* = 3 × 10^−4^; triangle in Figure 2). (E) KZFP-polymorphic zone with low maintenance cost (*s* = 5 × 10^−3^, *f* = 0.75, *c* = 3 × 10^−6^; square in Figure 2).

### KZFP-polymorphic zone

We next consider the KZFP-polymorphic zone where the KZFP suppressor allele does not fix but converges to an intermediate frequency (the green region in Figure 2A). In this zone, both the piRNA frequency, *p*_eq_, and the TE copy number, *n*_eq_, are generally low, particularly for KZFP with high efficacy ( *f*) and low cost (*c*).

The evolutionary dynamics of *x, p* and *n* in the KZFP-polymorphic zone are investigated in Figures 5D and E. With a large *c* in Figure 5D, *x* rises and settles at an intermediate level ( ∼0.4) while *p* and *n* drop more strongly than those in the KZFP-fixation zone with a smaller *f* . With a small *c* in Figure 5E, *x* overshoots transiently and then converges on ∼0.4 while *p* and *n* approach zero and occasionally exhibit sharp spikes.

This oscillation arises from a delayed negative feedback linking *x, p* and *n* through the KZFP suppression. As *x* increases, the average suppression efficacy 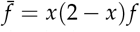 increases, leading to suppression of TE activity and a decline of *n* to near zero. When *n* becomes small, the fitness advantage of carriers of the KZFP suppressor allele over non-carriers vanishes, and the net selective advantage of KZFP, *σ* = *snu f* (1 −*p*)^2^ −*c*, falls below zero. Consequently, once *σ* < 0, *x* begins to decrease. The resulting weakening of suppression allows *n* to rebound, which in turn raises *σ* to positive values and drives *x* upward again. Because *p* responds to TE activity with a lag, via the first term in Equation (2), the phase delay between *x* and *p* produces an overshoot and a damped oscillation before the system stabilizes. In summary, in the KZFP-polymorphic zone, both systems coexist, but the high efficacy of KZFP drives the piRNA allele to a very low equilibrium frequency, sometimes near elimination.

## Discussion

Our study addresses the evolutionary puzzle of how multiple, seemingly redundant defense mechanisms against transposable elements can coexist within a single genome. By developing a three-component dynamical model of TEs, the piRNA system, and a newly introduced KZFP allele, we analytically characterized the conditions governing their interaction. Specifically, we derived the invasion threshold for a KZFP allele into an established TE-piRNA equilibrium (Equation (13)). Following a successful invasion, the KZFP allele can either spread to fixation or be maintained at an intermediate frequency, with the boundary determined by the condition based on Equation (22).

Our analysis reveals that, in the KZFP-fixation zone, piRNA and KZFP can coexist. In general, KZFP has an intermediate suppression efficacy and spreads to fixation due to the selective advantage conferred by its additional suppressive effect on TEs, whereas the selective pressure to maintain the piRNA system is reduced and its allele frequency declines but remains polymorphic. As a result, the cooperation of the two suppression systems effectively reduces TEs. When the suppression efficacy is too weak or the maintenance cost is too high, KZFP fails to establish. By contrast, when KZFP suppression is sufficiently strong, TE activity can be reduced to such a low level that the selective advantage of KZFP is weakened, allowing KZFP to be maintained at a polymorphic equilibrium.

However, this general pattern does not always hold. We find that TE copy number can paradoxically increase after KZFP introduction when selection against TEs is weak and/or the basal transposition rate is high. Under these conditions, TE proliferation is intrinsically strong, and the piRNA system plays a major role in TE suppression as its benefit outweighs its cost, whereas KZFP has only intermediate, rather than very strong, suppression efficacy (see above). This creates an interesting situation where KZFP partially suppresses TE activity but simultaneously weakens piRNA activity, because piRNA production requires TE insertions into piRNA clusters (i.e., piRNA tends to be more active when TEs are more active). In such cases, the total suppression capacity decreases because the KZFP is not strong enough to fully compensate for the weakened piRNA system. Consequently, a net increase in TE copy number occurs despite the addition of a new defense system. This paradoxical increase in TE number suggests that the allele-level selection does not necessarily maximize the population-level fitness. For carriers of KZFP, the allele confers a fitness advantage for individuals by reducing deleterious TEs (*σ* > 0), driving its spread. Yet as the KZFP allele becomes common, the piRNA allele decreases, allowing more TE copies to persist, potentially lowering mean population fitness compared with the pre-invasion equilibrium.

In the KZFP-polymorphic zone, the two defense systems do not coexist at appreciable frequencies in the long term. Although the KZFP allele is maintained at an intermediate frequency, its suppression efficacy is strong enough to reduce TE activity to a very low level. Because the piRNA system depends on ongoing TE transposition for its generation and maintenance, the piRNA-producing allele is effectively lost once TE activity is nearly eliminated. Moreover, when the maintenance cost *c* is very small, TE copy number *n* declines to almost zero. Under our deterministic infinite-population model, *n* never reaches zero exactly (Figure 5E), so that KZFP frequency *x* settles at a low value and does not disappear. However, this result may not hold in a finite population, where TEs are expected to go extinct stochastically due to random genetic drift. Once *n* reaches zero, the benefit of TE suppression vanishes and the net selective advantage of KZFP turns negative. In other words, KZFP-mediated suppression is so effective that it eliminates the TE threat entirely, making KZFP itself no longer necessary. Accordingly, cases such as Figure 5E should be interpreted as deterministic quasi-equilibria approached before stochastic TE extinction, rather than as stable long-term equilibria. This stochastic effect is indeed observed in our forward simulations described later.

Our analytical result for the invasion threshold for a KZFP allele (Equation (13)) produces a testable prediction: if a genome has a very large piRNA cluster fraction *π*, a KZFP allele should be less likely to invade and persist, because a large *π* reduces *n*^∗^ (Equation (10)) and the first term in *σ* (Equation (12)). Although a systematic phylogenetic analysis remains to be done, this prediction is qualitatively consistent with two well-studied species: *Drosophila melanogaster*, which lacks a KZFP system, has a relatively large piRNA cluster fraction (approximately 3.5% of its genome (Brennecke *et al*. 2007)), whereas humans, with a large KZFP repertoire, have a substantially smaller piRNA cluster fraction (roughly 0.4% (Rosenkranz *et al*. 2022)). These examples are illustrative rather than definitive, and a formal comparative analysis that regresses KZFP repertoire size against *π* across taxa—ideally using phylogenetic comparative methods and controlling for genome size, effective population size, and TE landscape—would provide a decisive test.

Our framework extends the established “trap model” of piRNA-mediated TE regulation (Kofler 2019; Tomar *et al*. 2023), in which TE insertions into piRNA clusters generate sequence-specific small RNAs that suppress the corresponding TE family. In the original trap model, the equilibrium TE copy number and piRNA-producing allele frequency are determined by the balance between transposition and purifying selection. In our model, we additionally include the benefit term for piRNA-producing alleles (Equation (2)), which arises because these alleles reduce the number of deleterious TE copies. This additional term decreases the predicted equilibrium TE copy number *n*^∗^ relative to the original trap-model formulation, whereas it does not change the pre-invasion equilibrium frequency *p*^∗^ (Equations (9) and (10)). This correction improves the quantitative agreement between the analytical predictions and the forward simulations (see below).

Introducing KZFP into this trap-model framework adds a second layer of sequence-specific suppression. KZFP-mediated regulation directly reduces the effective transposition rate from *u* to 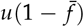 in both equations for the TE copy number and the piRNA frequency (Equations (1) and (2)). As a result, KZFP generally reduces the influx of both new TE insertions and new piRNA-producing alleles. However, when *s*/*u* is small, the reduction in piRNA-mediated suppression can lead to a paradoxical increase in equilibrium TE copy number after KZFP invasion (Equation (30)).

Our analysis is based on a simplified model that assumes a single TE family, but this simplification omits several features of real genomes. In reality, multiple TE families coexist and compete for shared resources such as insertion sites or replication machinery (Abrusén and Krambeck 2006; Venner *et al*. 2009; Lawlor and Ellison 2023). The expansion of one family can displace others and may alter their suppression dynamics in ways that our model does not capture. Additionally, our model does not incorporate the possibility of a continuous influx of novel TE families (Kofler *et al*. 2015; Scarpa *et al*. 2024). Given these factors that are ignored in our modeling framework, some of our results should be interpreted with caution. For example, when KZFP has strong suppression efficacy and a low maintenance cost, our model predicts near extinction of the resident TE family. The elimination of all TE copies would remove the selective pressure to maintain the costly KZFP and piRNA defense systems, leading to their eventual decay in frequency. However, this scenario is unlikely in real genomes: the loss of both defense systems would leave the genome vulnerable to invasion by new TE families. Future models that incorporate multiple TE families and multiple layers of defense systems will provide a more comprehensive understanding of these dynamics.

A further simplification of our model is the assumption of static sequences for both TEs and the two defense systems, which works for the analysis on short-term frequency dynamics. Over longer evolutionary timescales, however, sequence evolution is inevitable. TEs can accumulate mutations that allow them to escape recognition by piRNAs or KZFPs. In response, host genomes can evolve new defense variants—such as novel piRNA cluster insertions or mutations in the DNA-binding domains of KZFPs—that restore suppression against these escaping TEs. This reciprocal process is a molecular “arms race” between TEs and host defenses. Empirical observations support these recurrent cycles of conflict and adaptation at the sequence level. For example, TE families are often lineage-specific and show rapid turnover across species, meaning that new TE families emerge and expand while older ones become inactive or extinct. Similarly, the KZFP gene family exhibits rapid diversification, with frequent birth of new paralogs and loss of older ones (Thomas and Schneider 2011; Cosby *et al*. 2019; Bruno *et al*. 2019; Kosuge *et al*. 2024). Future models that incorporate sequence evolution would be able to explicitly simulate this arms race, providing deeper insights into the long-term co-evolutionary interaction between TEs and their hosts.

Another simplification used in our model is that TE insertions are assumed to occur uniformly across the genome. In reality, TE insertions can be non-random and such bias would alter the efficiency of TE insertion into piRNA clusters and thereby change the conditions for KZFP invasion and coexistence (Levy *et al*. 2010; Sultana *et al*. 2017; Pritam *et al*. 2025). If TEs exhibit an insertion bias that avoids piRNA clusters, the efficacy of the piRNA pathway would be substantially reduced. This would therefore make KZFP invasion more favorable. Conversely, a bias toward piRNA clusters would be expected to strengthen piRNA-mediated suppression and make KZFP invasion less favorable.

Although our main analysis focuses on the phylogenetically motivated scenario in which the piRNA system is already present before KZFP arises, the reverse or simultaneous order of the origin is also possible in principle. In numerical calculations initialized with KZFP regulation before the appearance of the piRNA-producing allele, or with simultaneous appearance of both defense systems, we found that the systems converged to the same equilibrium states as in the piRNA-first case, although the transient trajectories differed (not shown). Thus, the order of the origin mainly affects the short-term dynamics rather than the existence or location of the long-term equilibria in our deterministic analysis.

Furthermore, our model focuses exclusively on the interaction between the two key sequence-specific defense systems. In reality, these systems operate within a broader network of genome defenses that also includes sequence-independent but context-dependent silencing mechanisms, such as the HUSH complex in mammals, which senses long intronless nascent transcripts and reinforces H3K9me3-dependent repression (Seczynska *et al*. 2022; Nikolopoulos *et al*. 2025; Bloor *et al*. 2025). Incorporating such a mechanism into future models could provide a more comprehensive understanding of the multilayered TE suppression architecture.

Throughout this article, we use deterministic treatments that assume an infinite population size and free recombination among all loci. To assess whether our deterministic predictions remain robust when genetic drift and linkage disequilibrium are allowed, we performed forward individual-based simulations with finite population size, explicit chromosomes, and various levels of recombination. Our forward simulations further supported the robustness of the deterministic analysis. When selection against TE insertions was sufficiently strong, the three main outcomes predicted by the deterministic model—KZFP loss, KZFP fixation, and KZFP polymorphism—were all preserved. By contrast, when *s* was small, drift became more important, and stochastic fixation of the KZFP allele could occur even in regions where deterministic invasion was not predicted. Detailed results are given in Appendix E.

Our analytical model assumes that TE transposition events occur relatively independently, yielding a roughly Poisson-distributed TE copy number across the host population. For the parameter sets examined in our forward simulation, the variance in TE copy number was slightly above and close to the mean (not shown), suggesting that the results are not driven by extreme overdispersion. However, recent theoretical and empirical work suggests that TEs can sometimes exhibit “burst-like” transposition dynamics, leading to extreme overdispersion where the variance in TE copy number far exceeds the mean (Smith *et al*. 2022; Omole and Czuppon 2025). While our current framework does not explicitly model such transposition bursts, it is interesting to consider how extreme overdispersion might affect the coexistence of layered defense systems. Because positive selection on a suppressor allele like KZFP relies on the variance in TE copy number (i.e., the ability to associate with a genetic background with few TE copies), highly overdispersed TE distributions could theoretically enhance the efficiency of positive selection on suppressor alleles. This could effectively widen the parameter space for the successful invasion of novel defense mechanisms. Exploring the evolutionary dynamics of piRNA and KZFP systems under highly burst-like transposition regimes represents a fascinating avenue for future theoretical work.

Viewing the conflict between TEs and host suppression as a form of “genomic immunity” may provide a useful perspective for understanding why multiple layers of defense have evolved. From this perspective, piRNA-mediated defense resembles CRISPR-like memory-based immunity, in which sequence-specific suppression is established through the incorporation of foreign sequence information into a dedicated genomic locus (Marraffini 2015; Ophinni *et al*. 2019). By contrast, KZFP-mediated defense is more analogous to innate immunity, such as restriction factors or pattern-recognition receptors, because it relies on germline-encoded proteins to recognize specific molecular patterns and thereby provides constitutive defense (Pradeu *et al*. 2024). Our finding that a constitutive defense (KZFP) with only moderate suppression efficacy can inhibit the establishment of an activity-dependent defense (piRNA) may suggest an analogy in host immunity, where even moderately effective pathogen clearance by innate immunity could in principle limit the establishment of adaptive immune memory. Conversely, concepts from immune-evolution theory may help extend future TE-defense models (Mayer *et al*. 2016).

At the same time, an important difference between TE-defense systems and immune systems is that TEs are endogenous genomic parasites embedded in the host genome, so population-genetic processes such as linkage disequilibrium may play a more direct role than in standard host-pathogen models. Indeed, classical and recent population-genetic theories on the evolution of mutation and transposition modifiers have highlighted the critical role of linkage disequilibrium and recombination frequencies (Charlesworth and Langley 1986; Johnson 1999; Betancourt *et al*. 2024). Consistent with these established modifier theories, our individual-based simulations demonstrate that a negative association between KZFP allele dosage and TE copy number emerged, consistent with LD-mediated positive selection, without altering the outcomes predicted by our analytical model (see Appendix E).

**Appendices**

## APPENDIX A General model of KZFP allele frequency dynamics

In principle, KZFP efficacy may vary across many alleles. However, because the present analysis focuses on the invasion dynamics of a single suppressor allele against a null background, we use a minimal two-allele model in the main text. A more general efficacy-based formulation is provided here.

We consider a model that allows an infinite number of KZFP alleles. Allele *k* has suppression efficacy *f*_*k*_, ranging from 0 to 1, and frequency *x*_*k*_ . For simplicity, we do not explicitly model the sequence of the KZFP recognition motif; instead, we treat the efficacy *f*_*k*_ itself as a trait value. An allele with zero efficacy ( *f*_*k*_ = 0) is referred to as a null allele. The change in frequency of allele *k* over time is determined by natural selection and mutation:

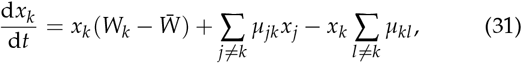

where ∑_*k*_ *x*_*k*_ = 1. *W*_*k*_ is the mean fitness of individuals carrying allele *k*, and 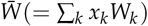 is the population mean fitness. *µ*_*jk*_ is the mutation rate from allele *j* to allele *k* per generation. Assuming that, in heterozygotes, the allele with the stronger suppression efficacy is dominant, the suppression efficacy of a diploid individual with genotype (*k, j*) is given by max( *f*_*k*_, *f*_*j*_).

Under Hardy-Weinberg equilibrium, the average suppression efficacy across the population is:

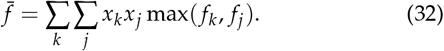

For analytical tractability, this general infinite-allele formulation is reduced to the two-allele model presented in the main text.

## APPENDIX B Jacobian Matrices at Equilibrium

To show whether the pre- or post-invasion equilibrium is stable or not, this appendix provides the Jacobian matrices for the two-variable (TE-piRNA) and three-variable (TE-piRNA-KZFP) systems, evaluated at their respective nontrivial equilibrium points. Substituting the equilibrium conditions to the Jacobian matrices allows for simplification of the matrix elements, clarifying the stability analysis.

### Jacobian for the TE-piRNA System (*x* = 0)

The two-variable system is described by Equations (7) and (8) with 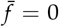. The nontrivial equilibrium point (*n*^∗^, *p*^∗^) satisfies the conditions:

i. *u*(1 − *p*^∗^)^2^ − *s* = 0

ii. 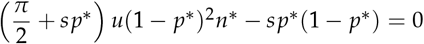.

Evaluating the Jacobian matrix at the equilibrium point (*n*^∗^, *p*^∗^), we have

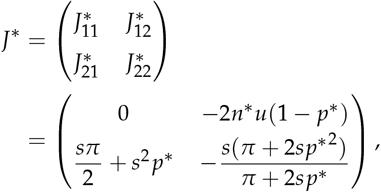

where the (1, 1) element of the Jacobian is 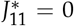 by the condition (i). Since 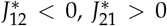, and 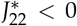 for 0 < *s* < *u* and *π* > 0, the determinant and trace are given by 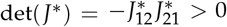 and 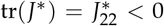, respectively. Therefore, the nontrivial TE-piRNA equilibrium is locally stable whenever it exists.

### Jacobian for the TE-piRNA-KZFP System (*x* > 0)

The full three-variable system is described by Equations (1),(2) and (6). Its nontrivial equilibrium (*n*_eq_, *p*_eq_, *x*_eq_) satisfies the conditions:

I.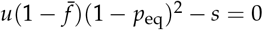

II.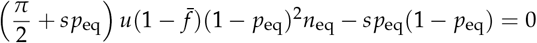

III. {*su f n*_eq_(1 − *p*_eq_)^2^ − *c*}*x*_eq_(1 − *x*_eq_)^2^ = 0.

Under these conditions, we have the Jacobian matrix:

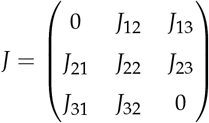

with

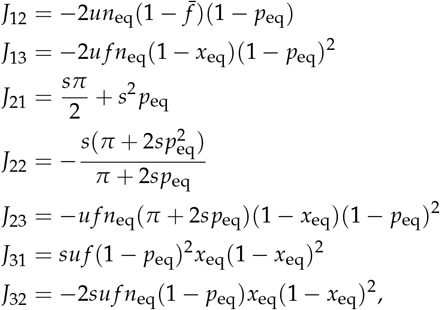

where *J*_11_ = *J*_33_ = 0 from the conditions (I) and (III).

Using the eigenvalues we obtained above, we examined whether the post-invasion equilibrium is stable or not in the ( *f, c*) plane. Figure A1 demonstrates that, regardless of whether KZFP fixes or remains polymorphic, the system consistently converges to a stable equilibrium. Each point in the ( *f, c*) plane is colored based on the eigenvalue structure of the Jacobian matrix *J* evaluated at equilibrium, confirming that the equilibrium is locally stable across all parameter sets examined. The blue region, located near the boundary between the KZFP-fixation and KZFP-polymorphic zones, corresponds to a stable node, where all variables converge directly to the equilibrium without oscillations. The surrounding cyan region indicates a stable focus, where convergence occurs with oscillations.

## APPENDIX C Quasi-equilibrium approximation—time-scale separation and validation

In the main text we treat the TE copy number *n* and the piRNA allele frequency *p* as fast variables and the KZFP allele frequency *x* as a slow variable. Here we quantify the assumption by comparing the relaxation rate of the (*n, p*) subsystem with the net selective advantage driving *x*.

For a fixed *x*, let (*n*_qs_, *p*_qs_) be the quasi-steady state obtained by solving 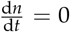 and 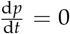 from Equations (1) and (2). Let *J*(*n, p* | *x*) be a 2 × 2 Jacobian of the (*n, p*) subsystem, and let *λ*_*i*_ denote its eigenvalues at (*n*_qs_, *p*_qs_). When the equilibrium is stable, the real part of the eigenvalues, denoted by ℜ*λ*_*i*_, is negative. Then we define the relaxation rate of fast variables (*n, p*) as *κ* and it holds

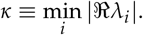

Under the quasi-equilibrium approximation, we have the net selective advantage of KZFP as

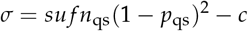

(cf. Equation (15)).

We quantify the timescale separation by a separation index, denoted by *ε*:

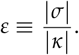

Heuristically, *ε* « 1 indicates that (*n, p*) relaxes to quasi-equilibrium much faster than *x* changes, justifying the quasi-equilibrium approximation at that *x*.

We also dynamically validate the quasi-equilibrium approximation by comparing the time courses of KZFP frequency *x* from the full three-dimensional (3D) model given by Equations (1), (2), and (6), with those from the one-dimensional (1D) reduced approximation described by Equation (14). Figure A2 shows the temporal dynamics of *x* for varying suppression efficacy *f* : dashed lines represent the full 3D system, and solid lines represent the 1D approximation. The trajectories from both models closely overlap, demonstrating that the reduced 1D model accurately captures the dynamics of the full system. This agreement holds even outside the fixation zone, where *f* is large ( *f* 0.9 in Figure A2), showing only mild phase lags when *κ* is small. These lags do not alter our conclusions and are not responsible for key phenomena such as the paradoxical increase in TE copy number.

## APPENDIX D Post-invasion equilibrium states in other parameter spaces

The overall patterns of the KZFP-fixation and KZFP-polymorphic zones remain robust across different parameter planes. Figure A3 shows the post-invasion equilibrium values of *x*_eq_, *p*_eq_, and *n*_eq_ plotted in the parameter space of basal transposition rate (*u*) and selection coefficient against TE insertions (*s*). Figure A4 shows the corresponding equilibrium values in the parameter space of piRNA cluster fraction (*π*) and selection coefficient against TE insertions (*s*).

## APPENDIX E Forward simulations

In the main text, we use a deterministic model to explore the evolutionary dynamics of two defense systems assuming an infinite-size population and free recombination between all loci, which successfully generates analytical expressions for insightful theoretical predictions. To examine whether these predictions from the deterministic model remain robust in a finite population with various levels of recombination, we performed forward individual-based simulations using Julia scripts (https://github.com/YusukeNabeka/TE-coexist).

In the simulations, populations consisted of *N* = 10,000 diploid individuals. Each individual had an explicit genome consisting of *L* = 10,000 possible TE insertion sites distributed across 30 chromosomes. One piRNA cluster was placed at the beginning of a single chromosome and occupied *πL* insertion sites. A single KZFP locus was placed at the end of a single chromosome outside the piRNA cluster. Each generation consisted of transposition, selection, and reproduction. During transposition, each TE copy produced a new insertion with an effective transposition rate determined by piRNA- and KZFP-mediated suppression. Individual fitness was then calculated from the deleterious effects of TE insertions and the maintenance cost of the KZFP allele. The next generation was sampled with replacement, with sampling probabilities proportional to individual fitness. Recombination was modeled using the Haldane mapping function (Haldane 1919). For each interval between adjacent insertion sites, the recombination fraction *r* was converted to a map distance 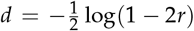. Recombination during gamete formation was then simulated along the resulting cumulative genetic map. We used *r* = 0.499 as an approximation to free recombination and *r* = 0.001 as a low-recombination case. The simulations were performed in two phases. First, TE–piRNA dynamics were simulated without KZFP until the system approached the pre-invasion equilibrium. Second, the KZFP suppressor allele was introduced into these populations at an initial frequency of 0.1, and the subsequent dynamics of TE copy number, piRNA allele frequency, and KZFP allele frequency were recorded.

A major difference in interpreting the results is that, in finite populations, new alleles are subjected to genetic drift. Thus, a KZFP allele could stochastically reach fixation even if its selective advantage is marginal, or conversely, it could be stochastically lost despite being selectively advantageous. To properly compare our stochastic simulations with the deterministic theoretical equilibria, we employed two key metrics. First, we evaluated the fraction of replicates in which the KZFP allele was established, that is, successfully invaded and persisted in the simulation. Second, for each parameter set, we recorded the conditional mean of the final values among the established replicates, thereby filtering out noise due to initial stochastic loss. Furthermore, in finite populations, TE copy number can also stochastically reach zero, whereas the deterministic model predicts that TE copy number approaches but never exactly reaches zero (see Figure 5E). In the simulations, once TEs are eliminated, KZFP loses its selective advantage and its frequency subsequently declines to zero slowly due to its maintenance cost. For this reason, in cases where TE extinction occurred, we recorded the corresponding values at the generation in which TE copy number reached zero, rather than capturing the subsequent, trivial decay of KZFP after TE loss.

Figure A5 shows simulation results in the ( *f, c*) parameter space. For each parameter set, simulations were replicated 100 times. The size of each plotted circle indicates the fraction of replicates in which the KZFP allele was established, while the color represents the conditional mean of the final values among the established replicates. Overall, the forward simulations confirmed that when *s* is large, the three major outcomes predicted by the deterministic model exist: KZFP loss, KZFP fixation, and KZFP polymorphism. Conversely, when *s* is small, the influence of genetic drift becomes more pronounced, allowing the KZFP allele to stochastically reach fixation even in parameter regions where invasion is deterministically predicted to fail. The fraction of replicates in which the KZFP allele was established was high when both the selection coefficient against TE insertions *s* and the KZFP suppression efficacy *f* were large, consistent with the expectation that the net selective advantage of the KZFP allele becomes larger under these conditions.

Unlike our deterministic model, which does not track linkage disequilibrium explicitly, our forward simulations with recombination can naturally generate linkage disequilibrium between TE insertion sites and the suppressor locus. To assess how linkage disequilibrium affects the behavior of the system, we measured the degree of linkage by calculating the covariance between suppressor allele dosage and TE copy number in the simulated populations, where allele dosage denotes the number of suppressor alleles carried by an individual (0, 1, or 2). Figure A6 displays representative time courses of TE copy number, piRNA frequency, KZFP frequency, and the covariance between suppressor allele dosage and TE copy number for representative parameter sets corresponding to KZFP loss, KZFP fixation, and KZFP polymorphism. During the periods in which KZFP frequency increased, the covariance between KZFP allele dosage and TE copy number became slightly negative as shown by olive colored lines in the lower panels of Figure A6B and C, indicating that genomes carrying KZFP tended to harbor fewer TE copies than genomes lacking KZFP. This pattern is consistent with the intuition that KZFP suppresses transposition in the genomes in which it is present, thereby generating linkage disequilibrium that contributes to positive selection on the KZFP allele. This pattern was observed under a wide range of recombination levels, including both very high recombination (at *r* = 0.499, Figure A6) and very low recombination (at *r* = 0.001, not shown). A negative covariance was also observed between piRNA allele dosage and TE copy number, as shown by the cyan lines in the lower panels of Figure A6. This indicates that genomes carrying piRNA-producing alleles also tended to have fewer TE copies. Thus, piRNA-producing alleles also receive a selective advantage through the reduction of deleterious insertions of TEs outside the piRNA cluster. This observation is consistent with the benefit term included in the analytical model (Equation (2)). Importantly, however, piRNA-producing alleles differ from KZFP alleles in that they also have a direct generation process of themselves through TE insertions into piRNA clusters. Therefore, the forward simulations support the interpretation that piRNA dynamics are shaped by both direct generation and positive selection, whereas KZFP dynamics depend primarily on positive selection on reducing TE copy number.

Finally, our main conclusions are also qualitatively robust to an alternative fitness function. In our main analysis, we adopted a simple log-linear fitness function. As an additional robustness check, we performed forward simulations under a log-quadratic fitness function that incorporates epistasis among TE copies, 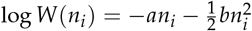, where *a* is the direct effect of a transposon insertion corresponding to *s* in our analytical model, and *b* represents the strength of synergistic epistasis among TEs (Dolgin and Charlesworth 2008; Roze 2023). Under this alternative fitness function, the patterns of KZFP loss, KZFP fixation and KZFP polymorphism are preserved (not shown). The equilibrium TE copy number was somewhat reduced and the invasion threshold shifted slightly, such that KZFP alleles with somewhat higher costs could invade and persist, because the benefit of suppressing TE copies became larger under the log-quadratic fitness function than under the log-linear fitness function.

## Data Availability

The authors state that all data necessary for confirming the conclusions presented in the article are represented fully within the article. The Python codes used for numerical analysis and generating the figures, and the Julia codes used for simulations are available at https://github.com/YusukeNabeka/TE-coexist.

## Acknowledgments

We sincerely thank the three anonymous reviewers for their valuable comments and suggestions. We also thank Ryuichi Kumata and Raiki Nakano for discussions on an early idea of this work, and Arnaud Le Rouzic for his helpful insights and feedback. The computing resources were provided by Human Genome Center (the Univ. of Tokyo).

---

**Figure A1.**
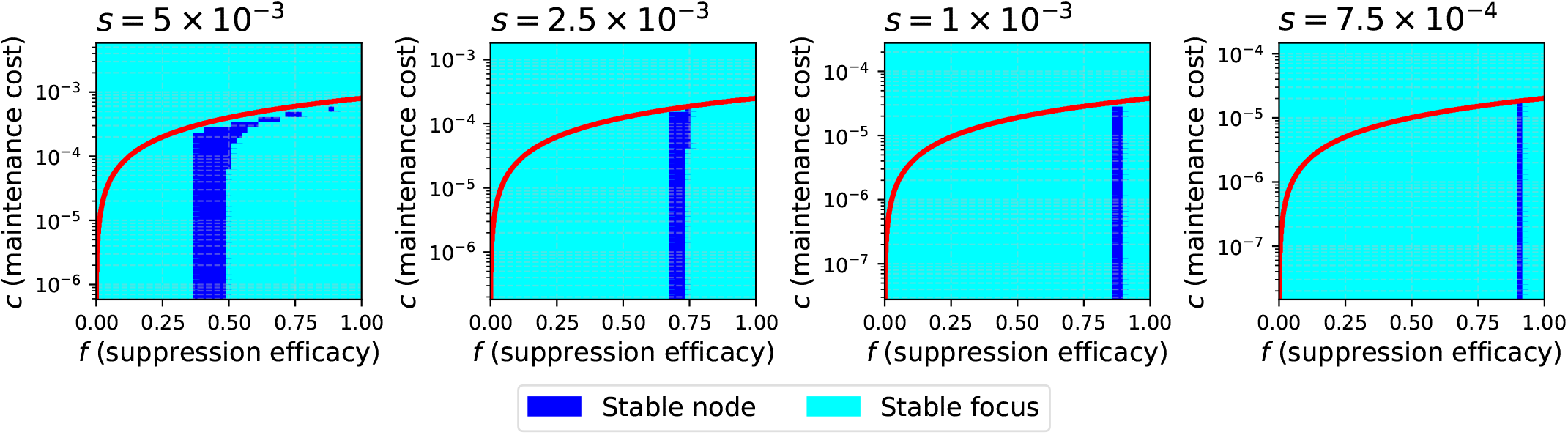
Local stability of the post-invasion equilibrium (*n*_eq_, *p*_eq_, *x*_eq_) in the ( *f, c*) plane. Parameter sets where the equilibrium is a stable node are shown in blue, and those where it is a stable focus are shown in cyan.

**Figure A2.**
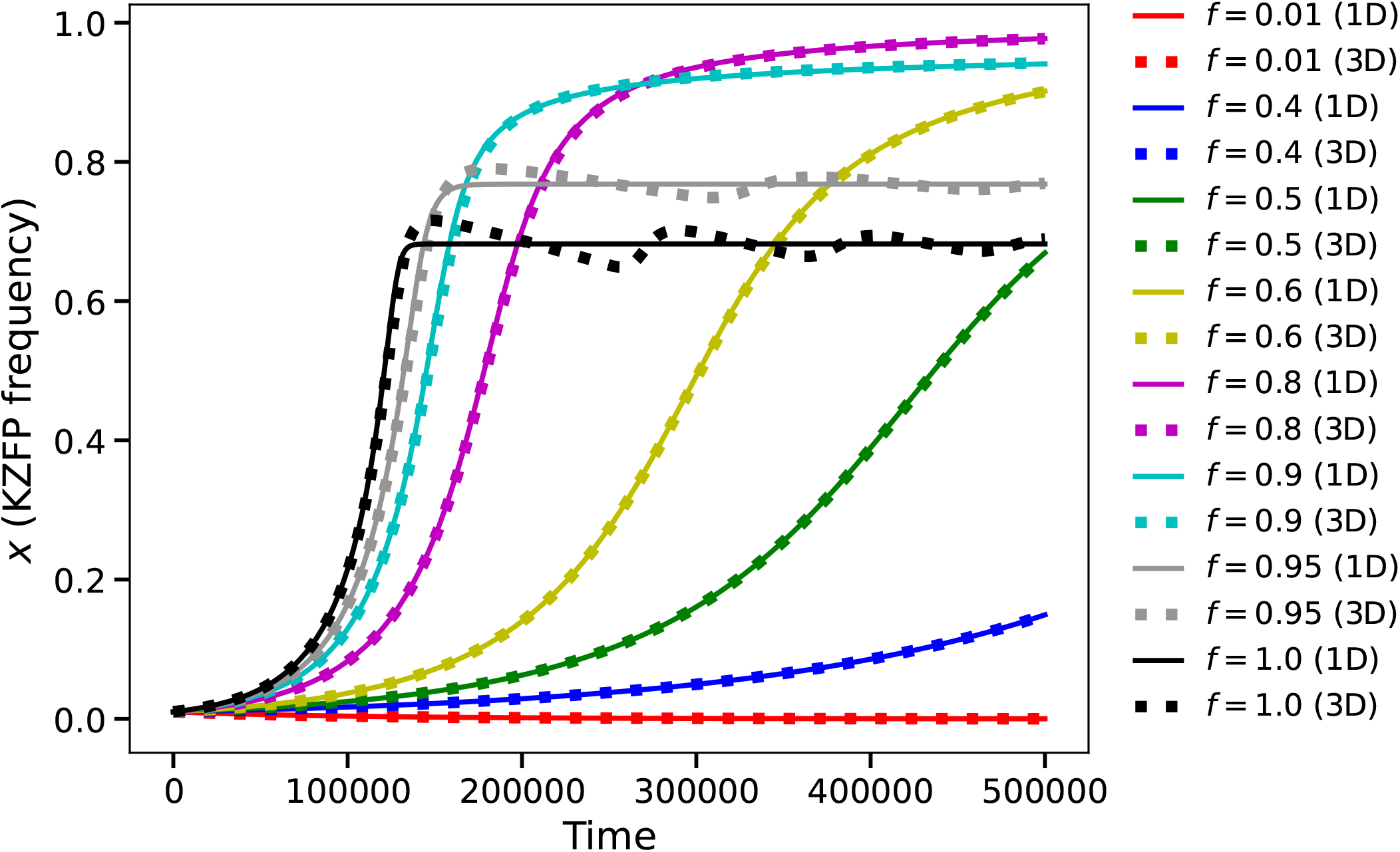
Validation of the quasi-equilibrium approximation. Time courses of KZFP frequency (*x*) from the full three-dimensional (3D) system (dashed lines) and the one-dimensional (1D) reduced model (solid lines) for representative values of suppression efficacy *f* . *u* = 0.01, *s* = 0.001, *π* = 0.01, and *c* = 1 × 10^−5^ are assumed. The reduced model closely tracks the full dynamics across all tested conditions. Minor deviations appear only at high efficacy ( *f* ≳ 0.9), where the time separation is less accurate.

**Figure A3.**
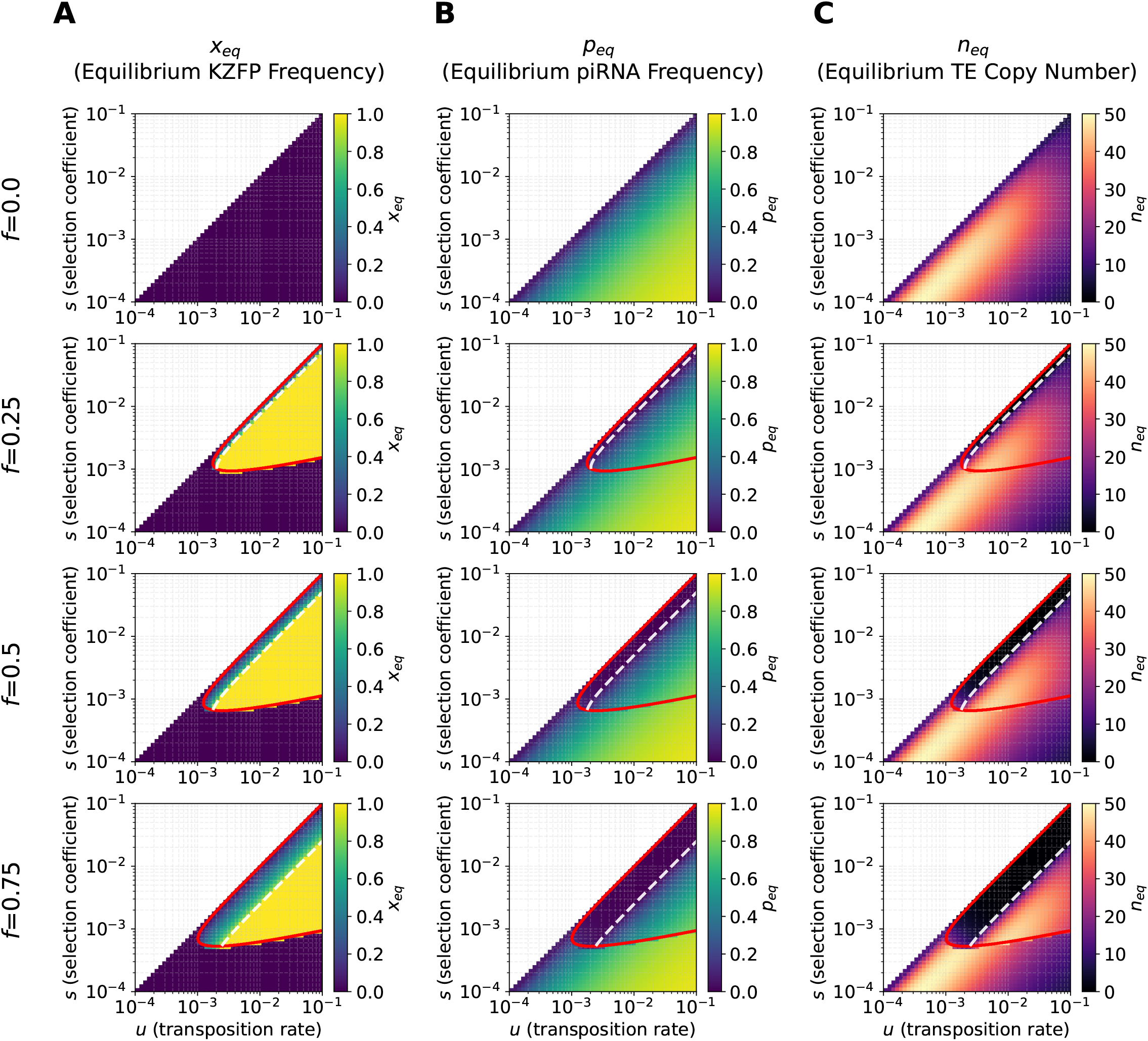
Post-invasion equilibrium values in the (*u, s*) parameter space. (A) Equilibrium frequency of the KZFP allele (*x*_eq_), (B) equilibrium frequency of the piRNA-producing allele (*p*_eq_), and (C) equilibrium TE copy number (*n*_eq_) after KZFP invasion plotted in the parameter space of basal transposition rate (*u*) and selection coefficient against TE insertions (*s*). Four different values of the suppression efficacy *f* are used ( *f* = 0.0, 0.25, 0.5 and 0.75), while *π* = 0.01 and *c* = 1 × 10^−5^ are fixed. The post-invasion equilibrium values are numerically computed by using the solve_ivp ODE solver in the scipy package (Virtanen *et al*. 2020). The color bars are to present the value of focal quantities in the post-invasion equilibrium. The red line shows the analytical invasion threshold from Equation (13), below which KZFP invasion is possible. The white dashed curve shows the upper boundary of the KZFP-fixation zone from Equation (22). The white area in the upper left corresponds to parameter combinations with *s* > *u*, where the TE-piRNA pre-invasion equilibrium does not exist because 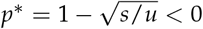 and *n*^∗^ < 0 (Equations (9) and (10)).

**Figure A4.**
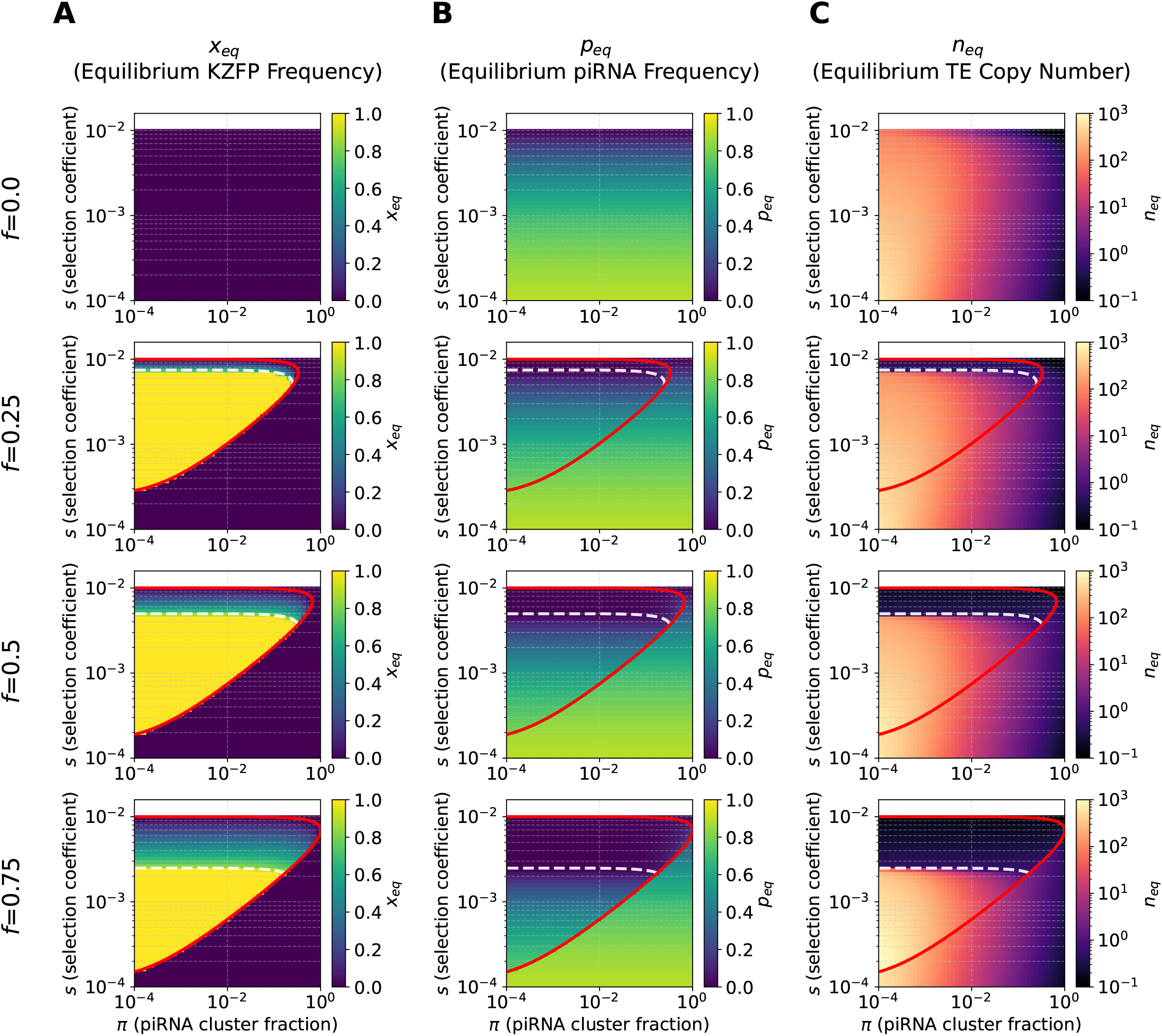
Post-invasion equilibrium values in the (*π, s*) parameter space. (A) Equilibrium frequency of the KZFP allele (*x*_eq_), (B) equilibrium frequency of the piRNA-producing allele (*p*_eq_), and (C) equilibrium TE copy number (*n*_eq_) after KZFP invasion plotted in the parameter space of piRNA cluster fraction (*π*) and selection coefficient against TE insertions (*s*). Four different values of the suppression efficacy *f* are used ( *f* = 0.0, 0.25, 0.5 and 0.75), while *u* = 0.01 and *c* = 1 × 10^−5^ are fixed. The post-invasion equilibrium values are numerically computed by using the solve_ivp ODE solver in the scipy package (Virtanen *et al*. 2020). The color bars are to present the value of focal quantities in the post-invasion equilibrium. The red curve shows the invasion threshold from Equation (13), below which KZFP invasion is possible. The white dashed curve shows the upper boundary of the KZFP-fixation zone from Equation (22). The white area in the upper corresponds to parameter combinations with *s* > *u*, where the TE-piRNA pre-invasion equilibrium does not exist because 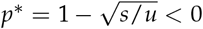 and *n*^∗^ < 0 (Equations (9) and (10)).

**Figure A5.**
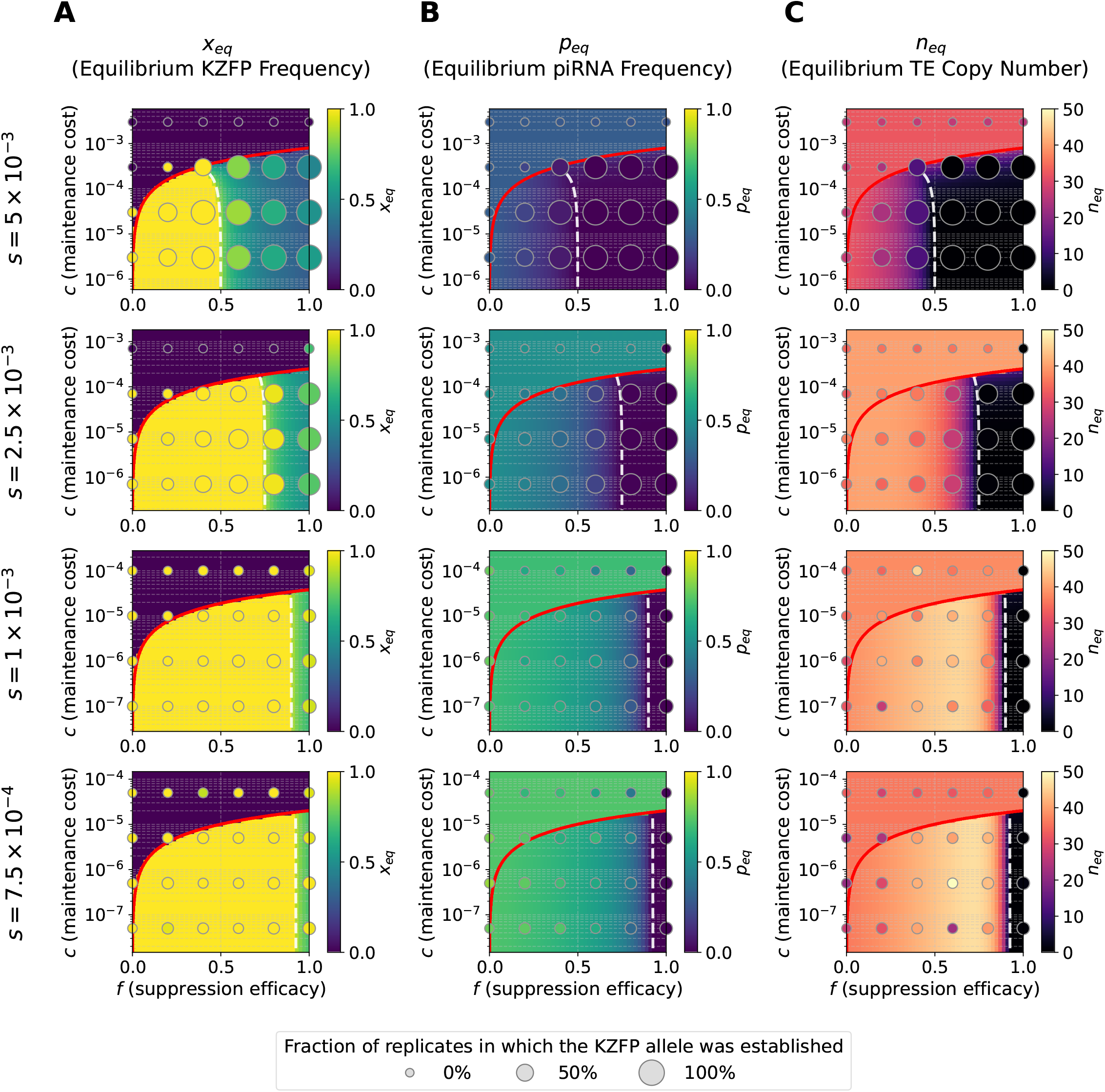
Results of the forward simulations overlaid on the post-invasion equilibrium colormaps in the ( *f, c*) parameter space of Figure 2. Panels show (A) KZFP frequency, (B) piRNA frequency, and (C) TE copy number. For each parameter set, simulations were replicated 100 times. The size of each circle indicates the fraction of replicates in which the KZFP allele was established, whereas the color indicates the conditional mean of the final values among the established replicates. The color bars apply both to the deterministic equilibrium values in the background colormaps and to the corresponding conditional mean values shown by the circles. Four different values of the selection coefficient against TE insertions, *s*, are shown, while *u* = 0.01, *π* = 0.01, and *r* = 0.499 are fixed. Other features are the same as in Figure 2.

**Figure A6.**
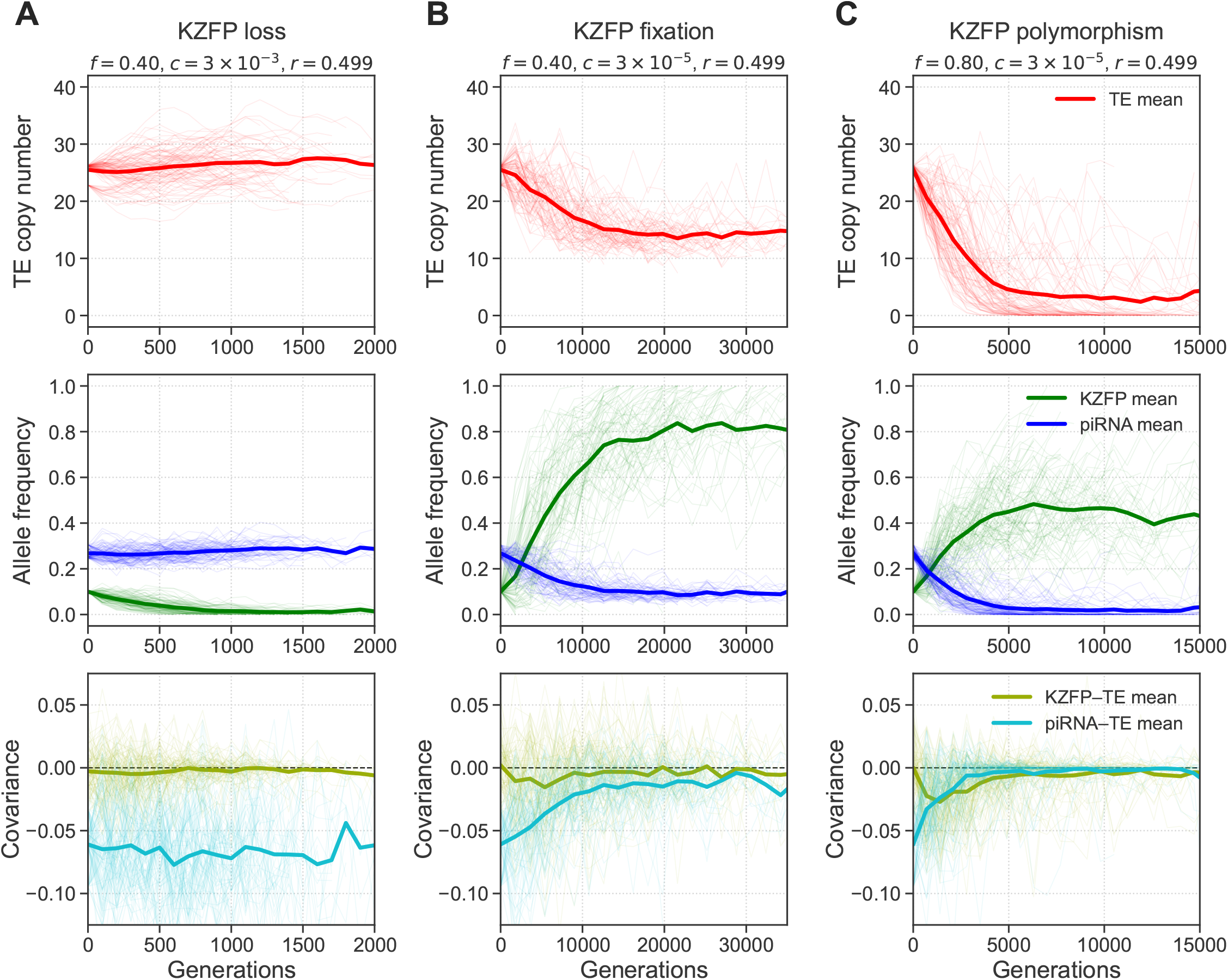
Trajectories from the forward simulations for three representative parameter sets from Figure A5: (A) KZFP loss (*s* = 5 × 10^−3^, *f* = 0.4, *c* = 3 × 10^−3^), (B) KZFP fixation (*s* = 5 × 10^−3^, *f* = 0.4, *c* = 3 × 10^−5^), and (C) KZFP polymorphism (*s* = 5 × 10^−3^, *f* = 0.8, *c* = 3 × 10^−5^). Fixed parameters were set to *u* = 0.01, *π* = 0.01 and *r* = 0.499. In each column, the upper panel shows TE copy number (red), the middle panel shows KZFP frequency (green) and piRNA frequency (blue), and the lower panel shows the covariance between suppressor allele dosage and TE copy number for KZFP (olive) and piRNA (cyan). Thin lines indicate individual replicates and thick lines indicate means of replicates.

## Literature cited

Abrusén G, Krambeck HJ. 2006. Competition may determine the diversity of transposable elements. Theoretical Population Biology. 70:364–375.

Almeida MV, Vernaz G, Putman AL, Miska EA. 2022. Taming transposable elements in vertebrates: from epigenetic silencing to domestication. Trends in Genetics. 38:529–553.

Betancourt AJ, Wei KHC, Huang Y, Lee YCG. 2024. Causes and consequences of varying transposable element activity: an evolutionary perspective. Annual review of genomics and human genetics. 25:1–25.

Bloor S, Wit N, Lehner PJ. 2025. Rna binding by periphilin plays an essential role in initiating silencing by the hush complex. Nucleic Acids Research. 53:gkae1165.

Brennecke J, Aravin AA, Stark A, Dus M, Kellis M, Sachidanandam R, Hannon GJ. 2007. Discrete Small RNA-Generating Loci as Master Regulators of Transposon Activity in Drosophila. Cell. 128:1089–1103.

Bruno M, Mahgoub M, Macfarlan TS. 2019. The Arms Race Between KRAB-Zinc Finger Proteins and Endogenous Retroelements and Its Impact on Mammals. Annual Review of Genetics. 53:393–416.

Charlesworth B, Charlesworth D. 1983. The population dynamics of transposable elements. Genetical Research. 42:1–27.

Charlesworth B, Langley C. 1986. The evolution of self-regulated transposition of transposable elements. Genetics. 112:359.

Cosby RL, Chang NC, Feschotte C. 2019. Host-transposon interactions: conflict, cooperation, and cooption. Genes & Development. 33:1098–1116.

Crow JF, Kimura M. 1970. An Introduction to Population Genetics Theory. Harper & Row. New York.

de Tribolet-Hardy J, Thorball CW, Forey R, Planet E, Duc J, Coudray A, Khubieh B, Offner S, Pulver C, Fellay J et al. 2023. Genetic features and genomic targets of human KRAB-zinc finger proteins. Genome Research. 33:1409–1423.

Dolgin ES, Charlesworth B. 2008. The effects of recombination rate on the distribution and abundance of transposable elements. Genetics. 178:2169–2177.

Ecco G, Imbeault M, Trono D. 2017. KRAB zinc finger proteins. Development. 144:2719–2729.

Emerson RO, Thomas JH. 2009. Adaptive evolution in zinc finger transcription factors. PLoS Genetics. 5.

Haldane JBS. 1919. The combination of linkage values, and the calculation of distances between the loci of linked factors. Journal of Genetics. 8:299–309.

Halic M, Moazed D. 2009. Transposon Silencing by piRNAs. Cell. 138:1058–1060.

Hanin G, Alsulaiti B, Costello KR, Tavares H, Takahashi N, Mikheeva LA, Freeman AK, Patel S, Jenkins B, Koulman A et al. 2026. Zfp57 is a regulator of postnatal growth and lifelong health. Nature Communications. .

Huntley S, Baggott DM, Hamilton AT, Tran-Gyamfi M, Yang S, Kim J, Gordon L, Branscomb E, Stubbs L. 2006. A comprehensive catalog of human KRAB-associated zinc finger genes: Insights into the evolutionary history of a large family of transcriptional repressors. Genome Research. 16:669–677.

Imbeault M, Helleboid PY, Trono D. 2017. KRAB zinc-finger proteins contribute to the evolution of gene regulatory networks. Nature. 543:550–554.

Iwasaki YW, Shoji K, Nakagwa S, Miyoshi T, Tomari Y. 2025. Transposon-host arms race: a saga of genome evolution. Trends in Genetics. 41:369–389.

Johnson T. 1999. Beneficial mutations, hitchhiking and the evolution of mutation rates in sexual populations. Genetics. 151:1621–1631.

Kazazian HH. 2004. Mobile Elements: Drivers of Genome Evolution. Science. 303:1626–1632.

Kofler R. 2019. Dynamics of Transposable Element Invasions with piRNA Clusters. Molecular Biology and Evolution. 36:1457–1472.

Kofler R, Hill T, Nolte V, Betancourt AJ, Schlötterer C. 2015. The recent invasion of natural drosophila simulans populations by the p-element. Proceedings of the National Academy of Sciences. 112:6659–6663.

Kosuge M, Ito J, Hamada M. 2024. Landscape of evolutionary arms races between transposable elements and KRAB-ZFP family. Scientific Reports. 14:23358.

Lawlor MA, Ellison CE. 2023. Evolutionary dynamics between transposable elements and their host genomes: mechanisms of suppression and escape. Current Opinion in Genetics & Development. 82:102092.

Levin HL, Moran JV. 2011. Dynamic interactions between transposable elements and their hosts. Nature Reviews Genetics 2011 12:9. 12:615–627.

Levy A, Schwartz S, Ast G. 2010. Large-scale discovery of insertion hotspots and preferential integration sites of human transposed elements. Nucleic acids research. 38:1515–1530.

Looman C, brink MA, Mark C, Hellman L. 2002. KRAB Zinc Finger Proteins: An Analysis of the Molecular Mechanisms Governing Their Increase in Numbers and Complexity During Evolution. Molecular Biology and Evolution. 19:2118–2130.

Luo S, Zhang H, Duan Y, Yao X, Clark AG, Lu J. 2020. The evolutionary arms race between transposable elements and pirnas in drosophila melanogaster. BMC Evolutionary Biology. 20:14.

Mani SR, Megosh H, Lin H. 2014. Piwi proteins are essential for early drosophila embryogenesis. Developmental biology. 385:340–349.

Marraffini LA. 2015. Crispr-cas immunity in prokaryotes. Nature. 526:55–61.

Mayer A, Mora T, Rivoire O, Walczak AM. 2016. Diversity of immune strategies explained by adaptation to pathogen statistics. Proceedings of the National Academy of Sciences. 113:8630–8635.

Najafabadi HS, Garton M, Weirauch MT, Mnaimneh S, Yang A, Kim PM, Hughes TR. 2017. Non-base-contacting residues enable kaleidoscopic evolution of metazoan C2H2 zinc finger DNA binding. Genome Biology. 18.

Nardelli J, Gibson T, Charnay P. 1992. Zinc finger-DNA recognition: analysis of base specificity by site-directed mutagenesis. Nucleic Acids Research. 20:4137–4144.

Nikolopoulos N, Oda Si, Prigozhin DM, Modis Y. 2025. Structure and methyl-lysine binding selectivity of the hush complex subunit mpp8. Journal of Molecular Biology. 437:168890.

Omole AD, Czuppon P. 2025. A population genetics model explaining overdispersion in active transposable elements. bioRxiv. https://www.biorxiv.org/content/early/2025/12/01/2025.11.27.691047.

Ophinni Y, Palatini U, Hayashi Y, Parrish NF. 2019. pirna-guided crispr-like immunity in eukaryotes. Trends in Immunology. 40:998–1010.

Ozata DM, Gainetdinov I, Zoch A, OCarroll D, Zamore PD. 2019. PIWI-interacting RNAs: small RNAs with big functions. Nature Reviews Genetics 2018 20:2. 20:89–108.

Pradeu T, Thomma BP, Girardin SE, Lemaitre B. 2024. The conceptual foundations of innate immunity: Taking stock 30 years later. Immunity. 57:613–631.

Pritam S, Scarpa A, Kofler R, Signor S. 2025. The impact of insertion bias into pirna clusters on the invasion of transposable elements. BMC biology. 23:258.

Rosenkranz D, Zischler H, Gebert D. 2022. piRNAclusterDB 2.0: update and expansion of the piRNA cluster database. Nucleic Acids Research. 50:D259–D264.

Rosspopoff O, Trono D. 2023. Take a walk on the KRAB side. Trends in Genetics. 39:844–857.

Roze D. 2023. Causes and consequences of linkage disequilibrium among transposable elements within eukaryotic genomes. Genetics. 224:iyad058.

Scarpa A, Pianezza R, Wierzbicki F, Kofler R. 2024. Genomes of historical specimens reveal multiple invasions of ltr retrotransposons in drosophila melanogaster during the 19th century. Proceedings of the National Academy of Sciences. 121:e2313866121.

Seczynska M, Bloor S, Cuesta SM, Lehner PJ. 2022. Genome surveillance by hush-mediated silencing of intronless mobile elements. Nature. 601:440–445.

Smith RD, Puzey JR, Conradi Smith GD. 2022. Population genetics of transposable element load: A mechanistic account of observed overdispersion. Plos one. 17:e0270839.

Srivastav SP, Feschotte C, Clark AG. 2024. Rapid evolution of piRNA clusters in the Drosophila melanogaster ovary. Genome Research. 34:711–724.

Sultana T, Zamborlini A, Cristofari G, Lesage P. 2017. Integration site selection by retroviruses and transposable elements in eukaryotes. Nature Reviews Genetics. 18:292–308.

Tadepally HD, Burger G, Aubry M. 2008. Evolution of C2H2-zinc finger genes and subfamilies in mammals: Species-specific duplication and loss of clusters, genes and effector domains. BMC Evolutionary Biology. 8.

Thomas JH, Schneider S. 2011. Coevolution of retroelements and tandem zinc finger genes. Genome Research. 21:1800–1812.

Tomar SS, Hua-Van A, Rouzic AL. 2023. A population genetics theory for piRNA-regulated transposable elements. Theoretical Population Biology. 150:1–13.

Venner S, Feschotte C, Biémont C. 2009. Dynamics of transposable elements: towards a community ecology of the genome. Trends in Genetics. 25:317–323.

Virtanen P, Gommers R, Oliphant TE, Haberland M, Reddy T, Cournapeau D, Burovski E, Peterson P, Weckesser W, Bright J et al. 2020. SciPy 1.0: fundamental algorithms for scientific computing in Python. Nature Methods. 17:261–272.

Wells JN, Chang NC, McCormick J, Coleman C, Ramos N, Jin B, Feschotte C. 2023. Transposable elements drive the evolution of metazoan zinc finger genes. Genome Research. 33:1325–1340.

Yang P, Wang Y, Macfarlan TS. 2017. The Role of KRAB-ZFPs in Transposable Element Repression and Mammalian Evolution.

Zhang Y, He F, Zhang Y, Dai Q, Li Q, Nan J, Miao R, Cheng B. 2022. Exploration of the regulatory relationship between KRAB-Zfp clusters and their target transposable elements via a gene editing strategy at the cluster specific linker-associated sequences by CRISPR-Cas9. Mobile DNA. 13:1–17.

